# Multiomics analyses decipher intricate changes in the cellular and metabolic landscape of steatotic livers upon dietary restriction and sleeve gastrectomy

**DOI:** 10.1101/2024.04.10.588826

**Authors:** Shuai Chen, Qinghe Zeng, Xiurong Cai, Jiaming Xue, Peng Song, Liming Tang, Christophe Klein, Frank Tacke, Adrien Guillot, Hanyang Liu

## Abstract

Metabolic dysfunction-associated steatotic liver disease (MASLD) is a chronic, progressive liver disease that encompasses a spectrum of steatosis, steatohepatitis (or MASH), and fibrosis. Evidence suggests that dietary restriction (DR) and sleeve gastrectomy (SG) can lead to remission of hepatic steatosis and inflammation through weight loss, but it is unclear whether these procedures induce distinct metabolic or immunological changes in MASLD livers. This study aims to elucidate the intricate hepatic changes following DR, SG or sham surgery in rats fed a high-fat diet as a model of obesity-related MASLD, in comparison to a clinical cohort of patients undergoing SG. Single-cell and single-nuclei transcriptome analysis, spatial metabolomics, and immunohistochemistry revealed the liver landscape, while circulating biomarkers were measured in serum samples. Artificial intelligence (AI)-assisted image analysis characterized the spatial distribution of hepatocytes, myeloid cells and lymphocytes. In patients and experimental MASLD rats, SG improved BMI, circulating liver injury biomarkers and triglyceride levels. Both DR and SG attenuated liver steatosis and fibrosis in rats. Metabolism-related genes (*Ppara*, *Cyp2e1* and *Cyp7a1*) were upregulated in hepatocytes upon DR and SG, while SG broadly upregulated lipid metabolism on cholangiocytes, monocytes, macrophages, and neutrophils. Furthermore, SG promoted myeloid cell accumulation in the liver not only ameliorating inflammation but activating liver repair processes. Regions with potent myeloid infiltration were marked with enhanced metabolic capacities upon SG. Additionally, a disruption of periportal hepatocyte functions was observed upon DR. In conclusion, this study indicates a dynamic cellular crosstalk in steatotic livers undergoing SG. Notably, PPARα- and gut-liver axis-related processes, and metabolically-active myeloid cell infiltration indicate intervention-related mechanisms supporting SG for the treatment of MASLD.

## Introduction

Obesity and overweight represent significant global health challenges, affecting approximately 2 billion people worldwide ^1^. Obesity-related complications encompass a spectrum of metabolic syndromes, including metabolic dysfunction-associated liver diseases (MASLD), metabolic dysfunction-associated steatohepatitis (MASH) and multi-organ disorders, contributing to elevated mortality rates among affected individuals ^2,3^. MASLD encompasses a range of chronic liver pathologies affecting more than a quarter of the global population, characterized predominantly by steatosis and fibrosis, with potential progression to cirrhosis and hepatocellular carcinoma and potent immune cell involvement at all disease stages ^4^. Despite considerable research efforts, pharmacotherapeutic targets for MASLD remain largely elusive, with only a limited number of candidates demonstrating efficacy. Only recently, resmetirom (a selective thyroid hormone receptor beta agonist) was approved by the Food and Drug Administration (FDA), USA as the first treatment of patients with MASH with moderate to advanced liver fibrosis, along with diet management and exercise ^5^.

Lifestyle modifications, such as dietary restriction (DR) and exercise, remain the mainstay of recommended therapeutic interventions in overweight and obese individuals with MASLD, with the aim of reducing body weight by at least 10% ^6^. However, the sustainability of long-term (hypocaloric) dietary management poses several challenges, particularly for individuals with severe obesity ^7^. Consequently, bariatric surgery (BS) has emerged as a viable option for obese patients, particularly when conventional nutritional and behavioral therapy fails to achieve desired therapeutic outcomes. In a randomized controlled clinical trial, BS-based interventions were superior to lifestyle intervention alone in weight loss as well as in improving histological features of MASLD such as resolution of MASH and improvement of liver fibrosis by at least 1 stage ^8^. Furthermore, observational studies demonstrated long-term benefits regarding cardiovascular and liver-related morbidity as well as mortality ^9^.

Among various surgical modalities, sleeve gastrectomy (SG) is the most frequently performed procedure, supported by robust evidence of long-term efficacy and safety ^10^. SG restricts nutritional intake by reducing the stomach volume and capacity, without reconstructing the gastrointestinal tract ^11^. Emerging studies have suggested that BS potentially mitigates lipid accumulation and halts MASLD progression ^12–14^. Although the primary consequence of SG is the restriction of food intake [akin to dietary restriction (DR)], SG generates remarkable influences systemic inflammation and gut microbiota composition ^15^. Recent advancements in omics technologies allow for sophisticated inquiries into liver diseases from diverse angles by profiling gene expression, protein function, metabolism, and microbiome activity ^16^. The latest study demonstrated the beneficial effects of SG on liver regenerative capacities based on mouse models ^17^. Simultaneously, our previous study demonstrated that BS exerts superior protective effects in the livers of patients with MASLD compared to DR, a finding associated with increased macrophage infiltration in BS ^13^. However, little has been known about metabolic and inflammatory alterations of steatotic livers after SG and DR interventions.

Here, we sought to compare the molecular and cellular consequences of DR and SG related to the amelioration of liver steatosis, using a rodent obesity-MASLD model. We implemented single-cell (sc)/single-nuclei (sn) transcriptome analysis to describe intracellular alterations and extracellular interactions, and spatial metabolomics with the assistance of artificial intelligence (AI)-based image analysis assists in deciphering metabolite production collating immune cell zonation. This study provides novel insights into multiomics alterations in steatotic livers undergoing DR and SG, which may hopefully facilitate understanding of mechanistic consequences of surgery-based therapies in MASLD.

## Results

### SG induces weight loss and ameliorates non-invasive biomarkers in human patients with MASLD

BMI and serological indexes were analyzed retrospectively in 18 patients who underwent SG (Fig. 1A). In comparison to baseline at one day pre-SG, significant declines of body mass index (BMI), glucose, postprandial blood glucose (PPG), aspartate aminotransferase (AST), alanine aminotransferase (ALT), gamma-glutamyltransferase (γ-GT), triglycerides (TG), high-density lipoprotein-cholesterol (HDL-C) and total bilirubin (TBIL) occurred at 6 months post-SG, while total bile acids (TBA) (p = 0.051) and fibroblast growth factor (FGF)-19 (*p* = 0.058) tended to increase (Fig. 1B-D, Table S2). This data hence revealed profound changes in varying processes, including inflammatory, metabolic and gut-derived endocrine signals. Intriguingly, male patients benefitted more from SG particularly on serological TG and low-density lipoprotein-cholesterol (LDL-C) levels than female patients (Table S2). Thus, and to further investigate the cellular mechanisms involved in the liver response to SG and compare with DR, healthy wild-type male rats were used to recapitulate interventions on an obesity-related MASLD model (Fig. 2A, Fig. S1). All rats gained weight after 12 weeks of high-fat diet (HFD) feeding, whereas Sham + DR and SG had a reduced weight gain in the 8 weeks following the intervention compared to HFD alone. Moreover, SG was the most efficient approach to halt weight gain (Fig. 2B). Serum analysis revealed that in comparison to Sham animals on HFD, Sham + DR rats had lower levels of LDL. SG significantly suppressed levels of TG, LDL and AST, but increased the levels of HDL, TBA and FGF-19. Additionally, SG drastically suppressed cholesterol levels (*p* = 0.06) but led to increased serum FGF-21 (*p* = 0.06) (Fig. 2C). HFD exaggerated hepatic lipid accumulation in comparison to rats fed a normal diet. Yet, both Sham + DR and SG intervention dramatically diminished hepatic lipid accumulation compared to the HFD-fed sham group (Fig. 2D and E). Simultaneously, fibrosis was increased after HFD, while it was significantly attenuated by both DR and SG (Fig. 2F and G). Results imply that both DR and SG effectively ameliorated obesity, liver steatosis and fibrosis, while SG was the most efficient intervention for improving systemic levels of lipid metabolism and bile acid (BA) production.

**Fig. 1.**
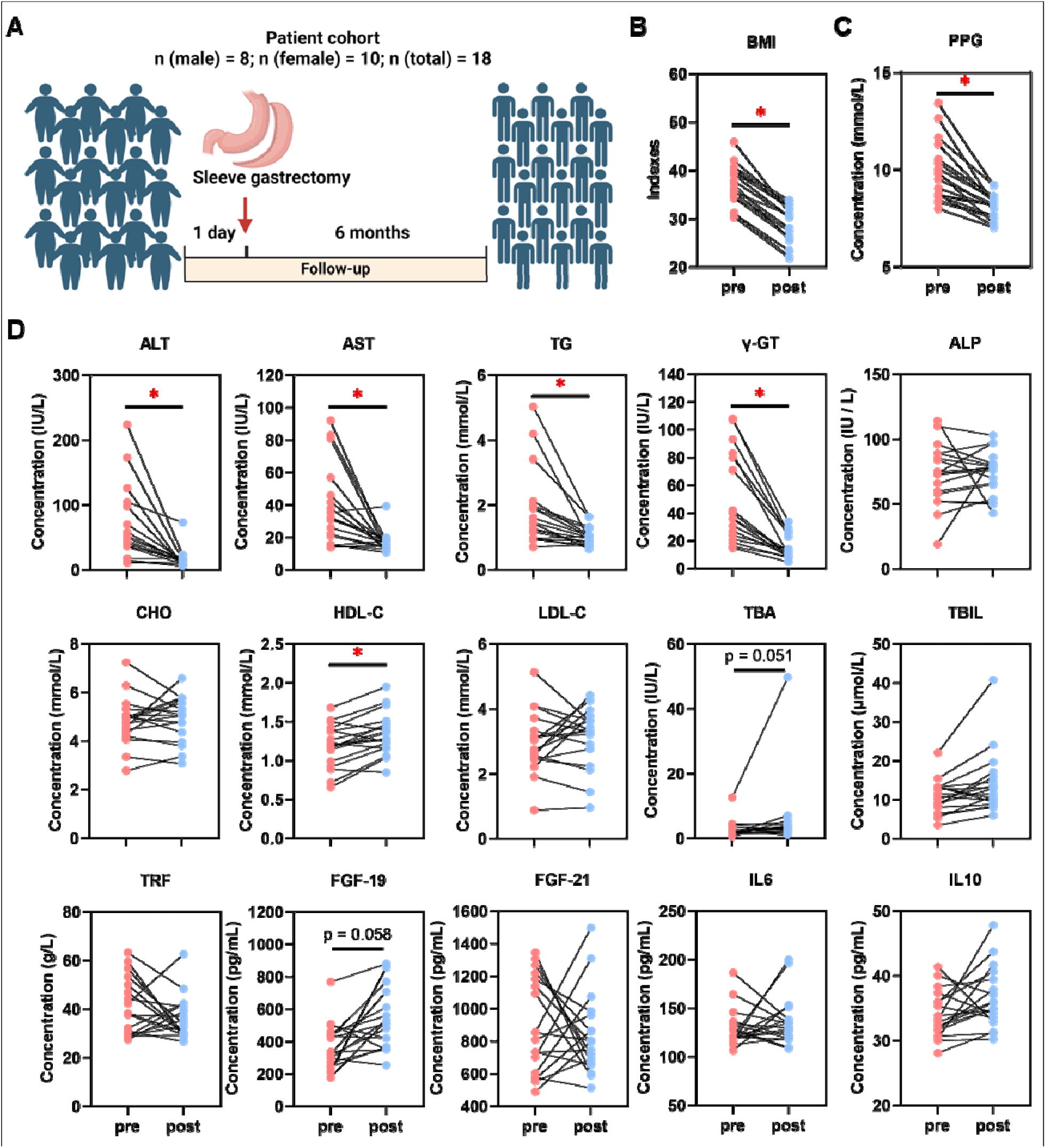
Sleeve gastrectomy induces weight loss and MASLD amelioration in patients. **(A)** Description of the patient cohort receiving SG. Comparisons of **(B)** BMI and **(C)** PPG, together with **(D)** ALT, AST, TG, γ-GT, ALP, CHO, HDL-C, LDL-C, TBA, TBIL, TRF, FGF-19, FGF-21, IL-6 and IL-10 in patients between one day pre-SG and 6 months post-SG. Abbreviations: SG: sleeve gastrectomy; BMI: body mass index; PPG: postprandial blood glucose; AST: aspartate aminotransferase; ALT: alanine aminotransferase; γ-GT: gamma-glutamyltransferase; TG, triglycerides; CHO, cholesterol; HDL-C, high-density lipoprotein cholesterol; LDL-C, low-density lipoprotein cholesterol; TBIL, total bilirubin; ALP, alkaline phosphatase; TBA, total bile acid. TRF: transferrin; IL: interleukin; FGF, fibroblast growth factor. The paired t-test was performed. ‘*’ represents ‘*p* < 0.05’ and statistical significance.

**Fig. 2.**
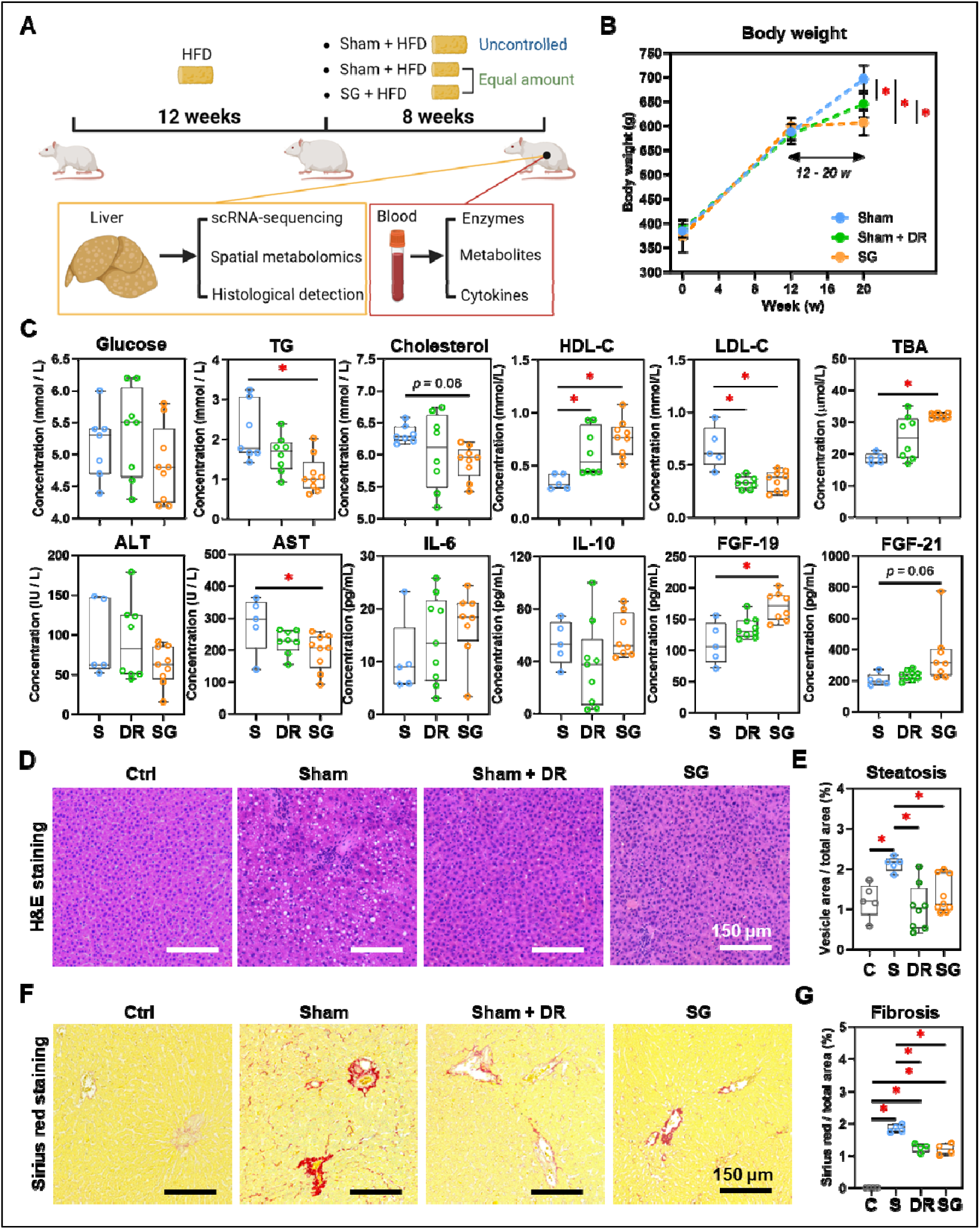
Sleeve gastrectomy and dietary restriction induce weight loss and MASLD amelioration in rat models. **(A)** The technical scheme of the rat MASLD model and surgical interventions. **(B)** Body weight was recorded for 20 weeks of modeling. **(C)** Serological indexes (glucose, triglycerides, cholesterol, HDL-C, LDL-C, TBA, ALT, AST, ALP, IL-6, IL-10, FGF-19 and FGF-21) were measured. **(D)** H&E-stained rat liver tissue and **(E)** quantification of steatosis (area of hepatocyte lipid vesicles) from Ctrl, Sham, Sham + DR and SG groups. **(F)** Sirius red-stained rat liver tissue and **(G)** quantification of fibrosis from Ctrl, Sham, Sham + DR and SG groups. Abbreviations: Sham/S: sham surgery; DR: dietary restriction; SG: sleeve gastrectomy; Ctrl: control fed a normal diet; HDL-C: density lipoprotein-cholesterol, LDL-C: low-density lipoprotein-cholesterol; TG: triglycerides; TBA: total bile acids, ALT: alanine transaminase; AST: aspartate aminotransferase; ALP: alkaline phosphatase; IL: interleukin; FGF: fibroblast growth factor. The one-way ANOVA test was performed. ‘*’ represents ‘*p* < 0.05’ and statistical significance.

### Sc/sn-transcriptome analysis evidences an altered hepatic cellular landscape after DR and SG

Rat liver samples [n=3 each per Sham (on HFD), Sham + DR and SG group] were analyzed using sc/snRNA-sequencing analysis, with an explicit classification of main hepatic and immune cell populations [including hepatocytes, cholangiocytes, hepatic stellate cells (HSC), endothelial cells, macrophages, monocytes, neutrophils, dendritic cells (DC), B lymphocytes, T lymphocytes and nature killer (NK) cells] (Fig. 3A, Fig. S2A and B). Reconstituted cell composition suggested higher numbers of NK cells, B cells, neutrophils and cholangiocytes leading to a lower proportion of hepatocytes and hepatic stellate cells upon Sham + DR (*vs.* Sham). In contrast, SG intervention led to increased proportions of macrophages and cholangiocytes but fewer monocytes and DCs (Fig. 3A). Metabolism-related marker expression (*Cyp2e1*, *Cyp7a1*, *Ppara*, *Nr1h4*, *Pnpla3* and *Gcg*) in liver steatosis was detected in hepatocytes and immune cells, and these pathways were transcriptionally regulated in Sham + DR and SG groups (*vs.* Sham) (Fig. 3B). Immunohistochemistry (IHC) evidenced that patatin like phospholipase domain containing 3 (PNPLA3) protein was significantly downregulated in Sham + DR and SG groups (*vs.* Sham), while peroxisome proliferator activated receptor (PPAR)α-expressing liver cells increased in the SG group (*vs.* Sham) (Fig. 3C and D), thereby confirming the profound regulation towards beneficial metabolic pathways in hepatocytes on a protein level after SG.

**Fig. 3.**
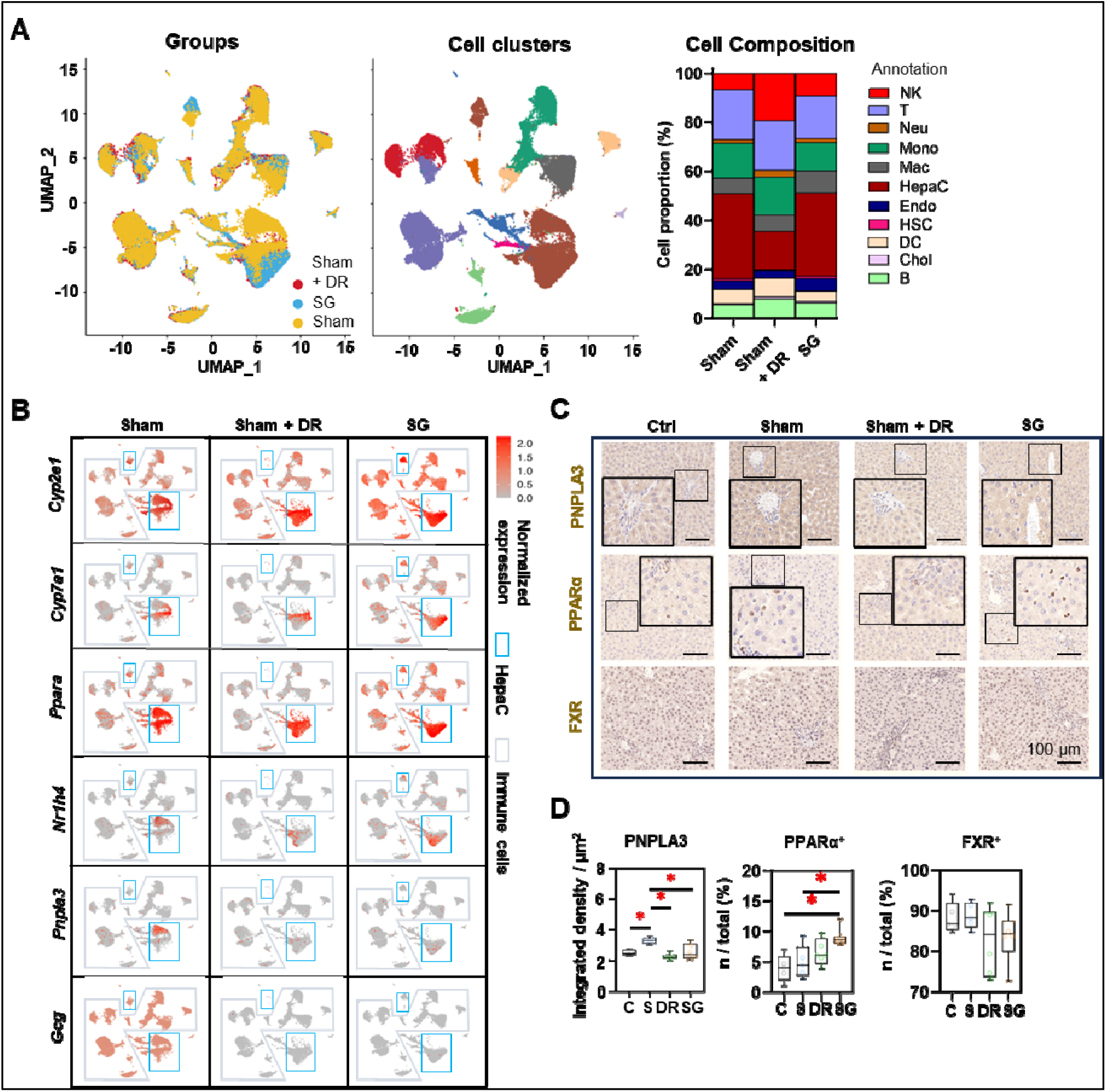
Sleeve gastrectomy improves hepatic metabolism according to sc/sn-transcriptome analysis and in-situ exploration. **(A)** UMAPs describe the distribution of B cells, cholangiocytes, DCs, endothelial cells, hepatic stellate cells, hepatocytes, macrophages, monocytes, neutrophils, T cells and NK cells. Histograms describing cell proportions. **(B)** Gene expressions of Cyp2e1, Cyp7a1, Ppara, Nr1h4, Pnpla3 and Gcg were illustrated on rat liver cells upon Sham, Sham + DR and SG. Protein levels of PNPLA3, PPARα and FXR were illustrated **(C)** and quantified **(D)** on rat liver tissue of Ctrl, Sham, Sham + DR and SG groups. Abbreviations: Ctrl: control fed a normal diet; Sham: sham surgery; DR: dietary restriction; SG: sleeve gastrectomy; PPAR: peroxisome proliferator-activated receptor; PNPLA: patatin-like phospholipase domain-containing protein; FXR: farnesoid X receptor; CK: cytokeratin; IL: interleukin; DC: dendritic cells; NK: natural killer cells; B: B lymphocytes; T: T lymphocytes; Neu: neutrophils; Mac: macrophages; Mono: monocytes: Endo: endothelial cells; HSC: hepatic stellate cells; Chol: cholangiocytes; HepaC: hepatocytes. The one-way ANOVA test was performed. ‘*’ represents ‘*p* < 0.05’ and statistical significance.

Furthermore, differentially expressed genes (DEGs) were screened in parenchymal (Fig. 4A) and immune cells (Fig. 4B) by comparing Sham + DR and SG to Sham groups. Transcriptomic influences of diverse cell types in the liver upon Sham + DR and SG were assessed based on cell proportions and significant DEG numbers. The distance of dots to value ‘0’ can be considered as the magnitude of influences. Data shows that hepatocytes and cholangiocytes were most influenced by SG, indicating a substantial impact of SG on metabolic functions of liver parenchymal cells, whereas endothelial cells were most influenced by Sham + DR (Fig. 4C), indicating ameliorated endothelial dysfunction as a consequence of DR. In accordance with the KEGG database, enriched signaling pathways were identified based on significantly upregulated DEGs. Up- and down-regulated DEGs at the bulk levels were compared between Sham, Sham + DR and SG groups, indicating enhanced metabolic processes (SG *vs.* Sham and SG *vs.* Sham + DR) and declined metabolic processes (Sham + DR *vs.* Sham) (Fig. S3B), which were in line with findings on liver samples from patients from our earlier study ^13^.

**Fig. 4.**
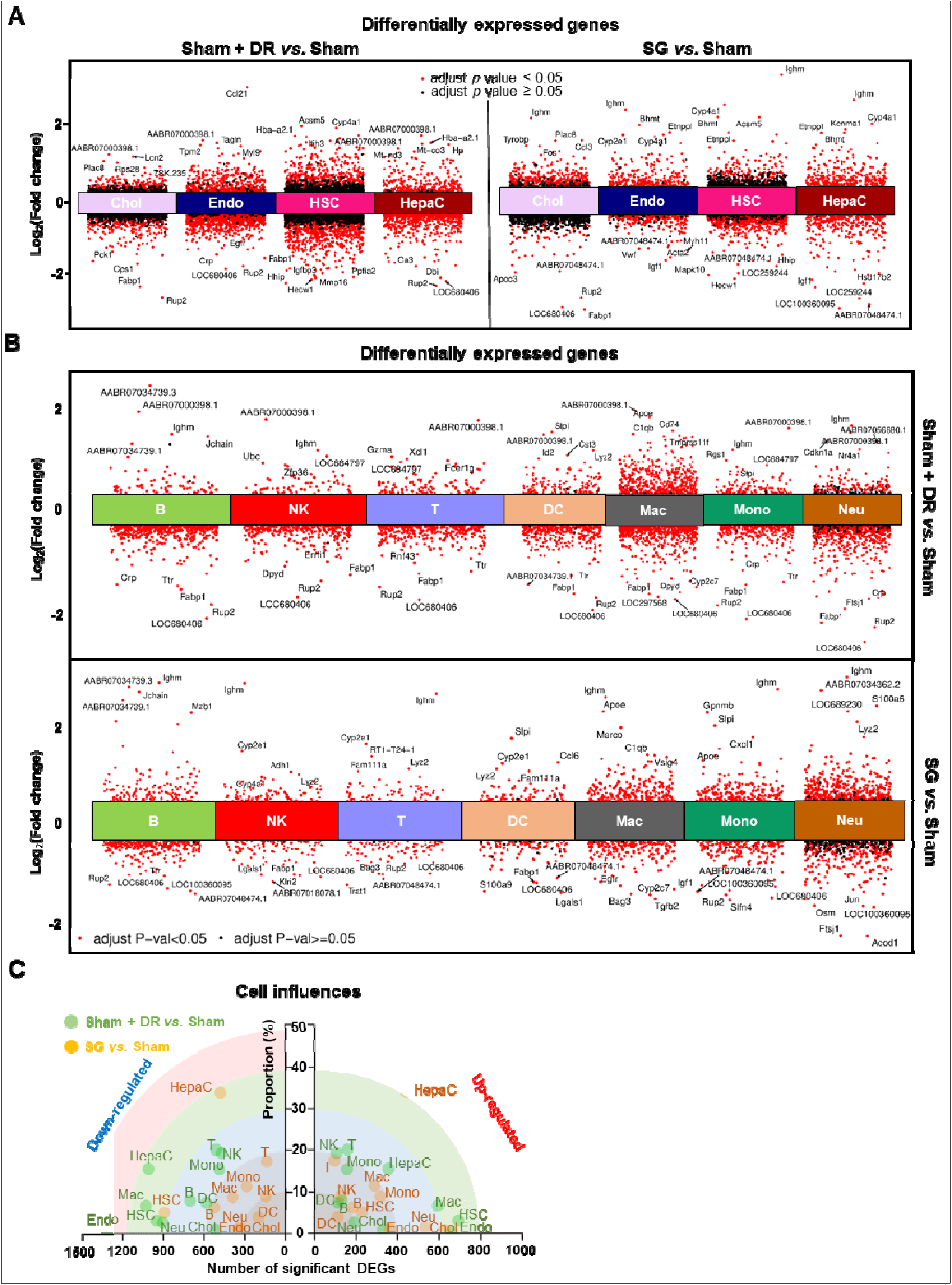
Sc/sn-transcriptome analysis illustrates differentially expressed genes in diverse liver cells. **(A & B)** Differentially expressed genes of hepatocytes, cholangiocytes, endothelial cells, hepatic stellate cells, B, T, NK, DCs, macrophages, monocytes and neutrophils (Sham + DR *vs.* Sham and SG *vs.* Sham) were illustrated in volcano plots. **(C)** Transcriptomic influences of hepatocytes, cholangiocytes, endothelial cells, hepatic stellate cells, B, T, NK, DCs, macrophages, monocytes and neutrophils in the liver were assessed with DEG numbers and cell proportion of total cells Abbreviations: Sham: sham surgery; DR: dietary restriction; SG: sleeve gastrectomy; DC: dendritic cells; NK: natural killers; B: B lymphocytes; T: T lymphocytes; Neu: neutrophils; Mac: macrophages; Mono: monocytes: Endo: endothelial cells; HSC: hepatic stellate cells; Chol: cholangiocytes; HepaC: hepatocytes; DEG: differentially expressed gene. The one-way ANOVA test was performed. ‘Adjust *p* < 0.05’ represents statistical significance.

Next, regulated signaling pathways were compared between the groups (Sham + DR *vs.* Sham, SG *vs.* Sham, and SG *vs.* Sham + DR) for each cell population (Fig. 5A). Fatty acid degradation-related gene expression was increased not only in hepatocytes (Sham + DR *vs.* Sham, SG *vs.* Sham, SG *vs.* Sham + DR) but also in cholangiocytes and endothelial cells (SG *vs.* Sham). Furthermore, PPAR signaling pathways, fatty acid degradation and cholesterol metabolism were more pronounced (not only in hepatocytes) by both Sham + DR and SG interventions (*vs.* Sham). Interestingly, in comparison to Sham + DR, the PPAR signaling pathway was promoted in cholangiocytes, endothelial cells and myeloid cell populations (monocytes, macrophages, DCs and neutrophils). Of note, butanoate metabolism was enhanced in hepatocytes upon both Sham + DR and SG (Fig. 5A). In addition, cellular interactions (total and ligand-receptor) were elucidated between each gradual pair of cell types among all three groups using the CellChat algorithm (Fig. S4A and B). Key signaling pathways that enriched according to total cellular interactions were described (Fig. S5A). In contrast to Sham, both Sham + DR and SG elevated ligand reception of neutrophils from immune cells, while SG elevated ligand release from cholangiocytes to other cells. Of note, in contrast to Sham + DR groups, SG promoted interaction levels of non-hepatocyte parenchymal cells (cholangiocytes, HSCs and endothelial cells) as well as immune cells (mainly for macrophages and neutrophils) (Fig. 5B). Further dissecting cholangiocyte-associated cellular interactions, significant signaling pathways from cholangiocytes to monocytes, macrophages, neutrophils, B lymphocytes, T lymphocytes and NK cells, as well as from monocytes, macrophages, DCs, hepatocytes, T lymphocytes and NK cells to cholangiocytes were illustrated (Fig. S5B and C).

**Fig. 5.**
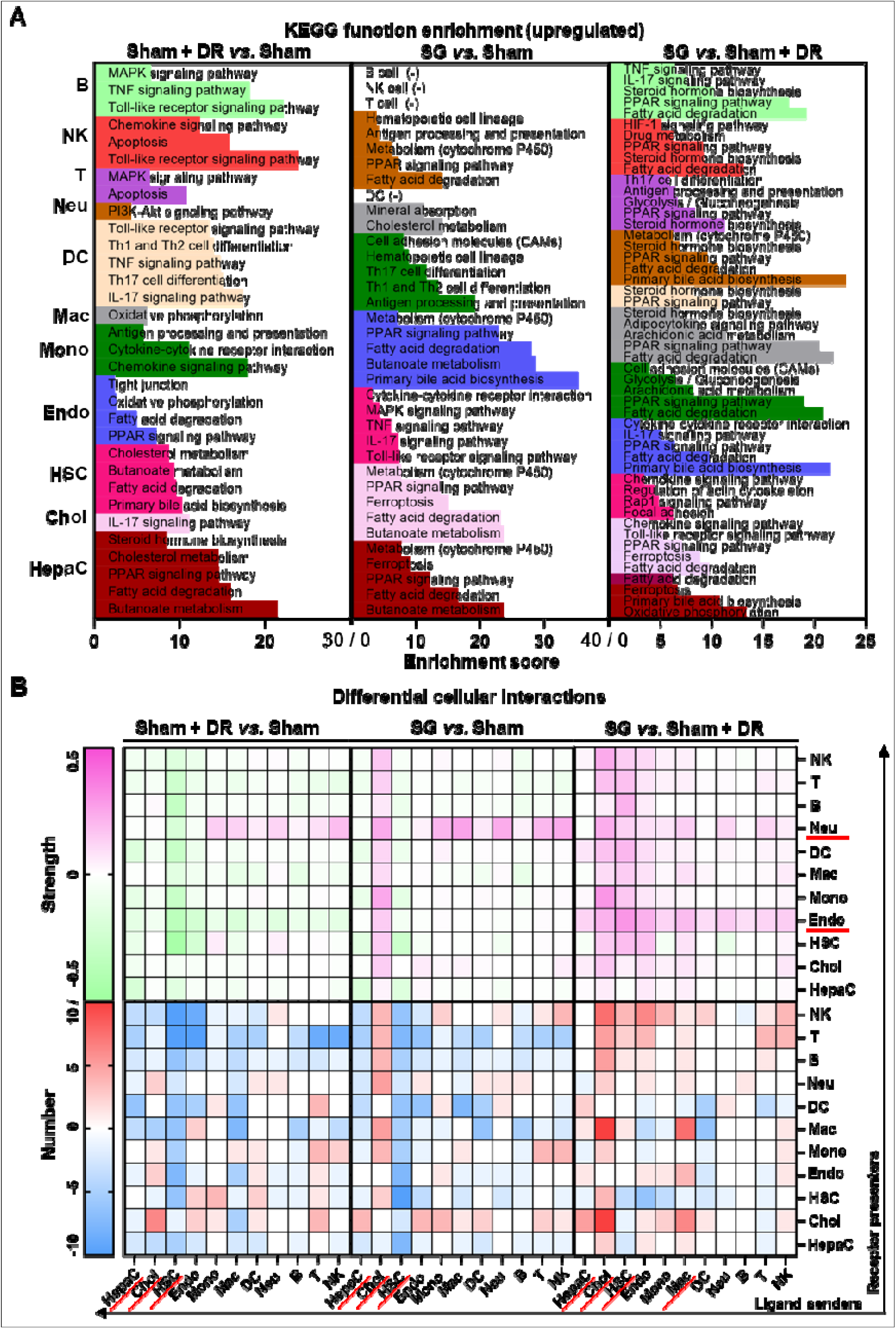
Sc/sn-transcriptome analysis illustrates up-regulated enriched pathways and cellular interactions in diverse liver cells. Significantly up-regulated **(A)** KEGG signaling pathways and **(B)** cellular interactions (numbers and strength) in hepatocytes, cholangiocytes, endothelial cells, hepatic stellate cells, B, T, NK, DCs, macrophages, monocytes and neutrophils were illustrated (Sham + DR *vs.* Sham and SG *vs.* Sham). Abbreviations: Sham: sham surgery; DR: dietary restriction; SG: sleeve gastrectomy; HepaC: hepatocytes; Chol: cholangiocytes; HSC: hepatic stellate cells; Endo: endothelial cells; Mono: monocytes; Mac: macrophages; DC: dendritic cells; Neu: neutrophils; NK: natural killer cells; B: B lymphocytes; T: T lymphocytes; KEGG: Kyoto encyclopedia of genes and genomes. The one-way ANOVA test was performed. ‘Adjust *p* < 0.05’ represents statistical significance.

Taken together, both Sham + DR and SG improve liver lipid metabolism (remarkably PPAR-associated processes) in steatotic livers. In particular, SG triggers metabolic enhancement broadly in liver cells, including hepatocytes, cholangiocytes, endothelial cells and myeloid cells. Additionally, cholangiocytes appear to actively interact with multiple liver cell types upon SG.

### SG promotes liver metabolic capacities by accumulating myeloid cell population

To explore the metabolic landscape of the liver, three representative liver samples were freshly acquired from the Sham, Sham + DR and SG groups and then analyzed using an ambient mass spectrometry imaging (MSI) metabolomics approach (Fig. 6A). The spectrum and distribution of total metabolites (with positive and negative ions) were illustrated (Fig. S6A – E). Histological features of liver samples together with spatially distributed metabolite abundance were displayed, indicating strongly reduced production of total metabolites in steatotic livers undergoing DR or SG (Fig. 6B). Function enrichment analysis based on the total abundance of differentially upregulated metabolites illustrated that both Sham + DR and SG enhanced lipid metabolism in steatotic livers in contrast to Sham groups, potentially attenuating steatotic stress in liver. It is worth noting that SG significantly enhanced the capacities of linoleic acid metabolism, fat digestion and absorption, and cholesterol metabolism compared to Sham + DR (Fig. 6C).

**Fig. 6.**
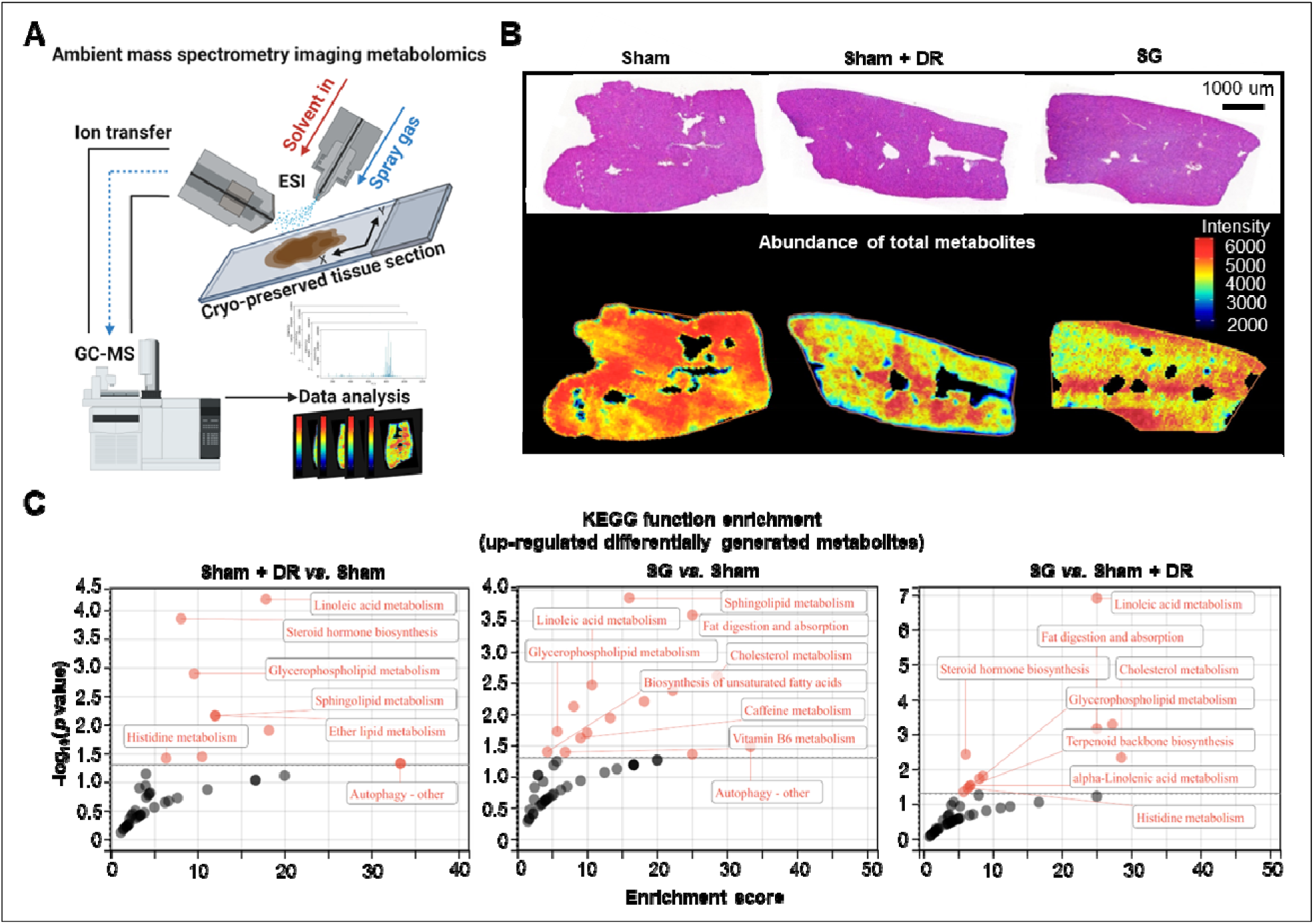
Sleeve gastrectomy and dietary restriction enhance metabolic capacities in steatotic livers. **(A)** Technical scheme of the ambient mass spectrometry imaging metabolomics method. **(B)** H&E-stained rat liver sample section was mapped with the abundance of total metabolites in Sham, Sham + DR and SG groups. **(C)** Function enrichment of upregulated differentially generated metabolites (Sham + DR *vs.* Sham, SG *vs.* Sham and SG *vs.* Sham + DR) was analyzed based on the KEGG database. Abbreviations: Sham: sham surgery; DR: dietary restriction; SG: sleeve gastrectomy; KEGG: Kyoto encyclopedia of genes and genomes. The one-way ANOVA test was performed. ‘Adjust *p* < 0.05’ represents statistical significance.

AI-based pathological exploration has evolved as a promising and precise tool for MASLD investigations ^18^. Hence in this study, a deep-learning model was established based on cell annotations (hepatocytes, lymphocytes and myeloid cells) that were supervised by liver pathology experts (Fig. 7A). Accordingly, numbers of hepatocytes, lymphocytes and myeloid cells were assessed on a total of 18 rat liver samples. Importantly, hepatocyte proportions (70 – 80%) of total liver cells were not significantly altered by different interventions (Fig. 7B). Lymphocyte proportions were significantly increased in Sham with HFD (1.5 – 6.1%) *vs.* healthy control (0.05 −2.1%) but declined upon Sham + DR (0.7 – 2.0%) and SG (0.5 – 2.2%) (*vs.* Sham) (Fig. 7B). Simultaneously, SG exacerbated the infiltration of myeloid cells (*vs.* healthy control, *vs.* Sham + DR) (Fig. 7B). In comparison, the numbers of lymphocytes and myeloid cells were assessed based on the sc/sn RNA-seq dataset. Myeloid cell proportions tended to increase by SG (Fig. S7A and B). Besides, the cell distribution of lymphocytes and myeloid cells was depicted to correspondingly map the metabolites in diverse cell regions (Fig. 7C). The abundance of total metabolites between regions with high or low myeloid cell infiltration was compared (high-/low-infiltration region selection displayed in Fig. 7C). SG remarkably increased metabolite abundance in regions with high myeloid cell infiltration (Fig. 7D). In addition, upregulated metabolites in regions rich in myeloid cells significantly refer to processes of autophagy, together with the metabolism of choline, linoleic acids, cholesterol and glycerophospholipids (Fig. 7E). Given the diverse phenotypic functions of macrophages in liver, a list of markers (*Trem2*, *Spp1*, *Gpnmb*, *Mmp9*, *Mmp13*, *Plin2*, *Ppara*, *Pparg*, *Cd207*, *Cx3cr1*, *Clec4f*, *Nos2*, *Cd80*, *Nlrp3*, *Il1b*, *Tnfa*, *Cd163*, *Mrc1*, *Il10* and *Tgfb1*) was selected from the sc/sn RNA-seq dataset to assess phenotypes of scar-associated macrophage (SAM), lipid-associated macrophage (LAM), liver capsular macrophage (LCM), Kupffer cell (KC), pro- and anti-inflammatory macrophages ^19–21^. Comparing macrophage clusters (SG *vs.* Sham), markers of SAM, LCM and KC were upregulated (SG *vs.* Sham), whereas pro-inflammatory macrophage markers were downregulated (SG *vs.* Sham and Sham + DR *vs.* Sham) (Fig. 7F, Fig. S7C). Simultaneously, SG promoted classical phenotypes of monocytes bur suppressed neutrophil maturation. In contrast, DR tended to promote conventional DC phenotype and neutrophil maturation (Fig. 7F, Fig. S7C). Taken together, SG and DR enhanced lipid metabolism in the liver, while SG promoted myeloid cell accumulation in the liver, facilitating metabolic improvement and liver repair processes.

**Fig. 7.**
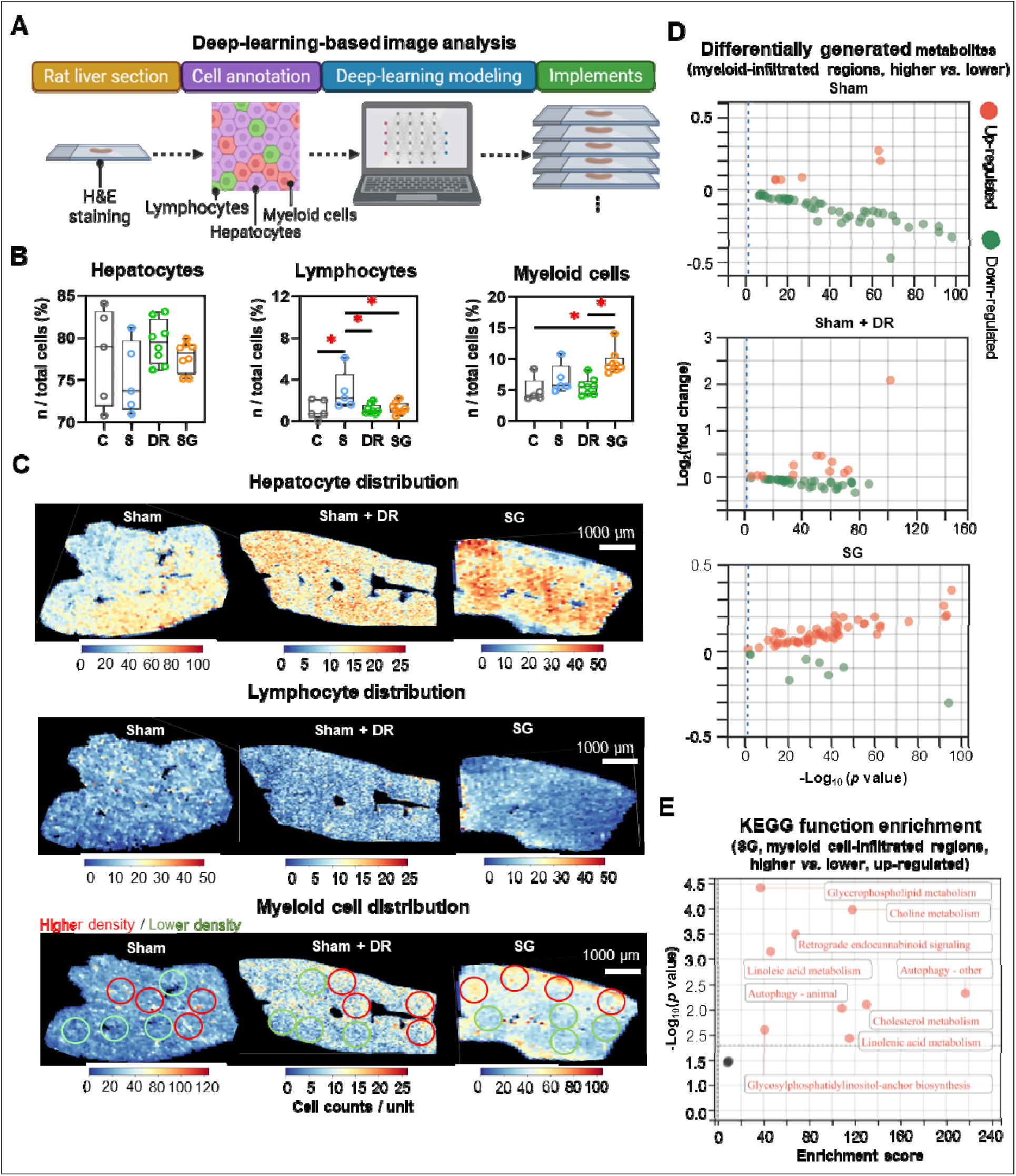
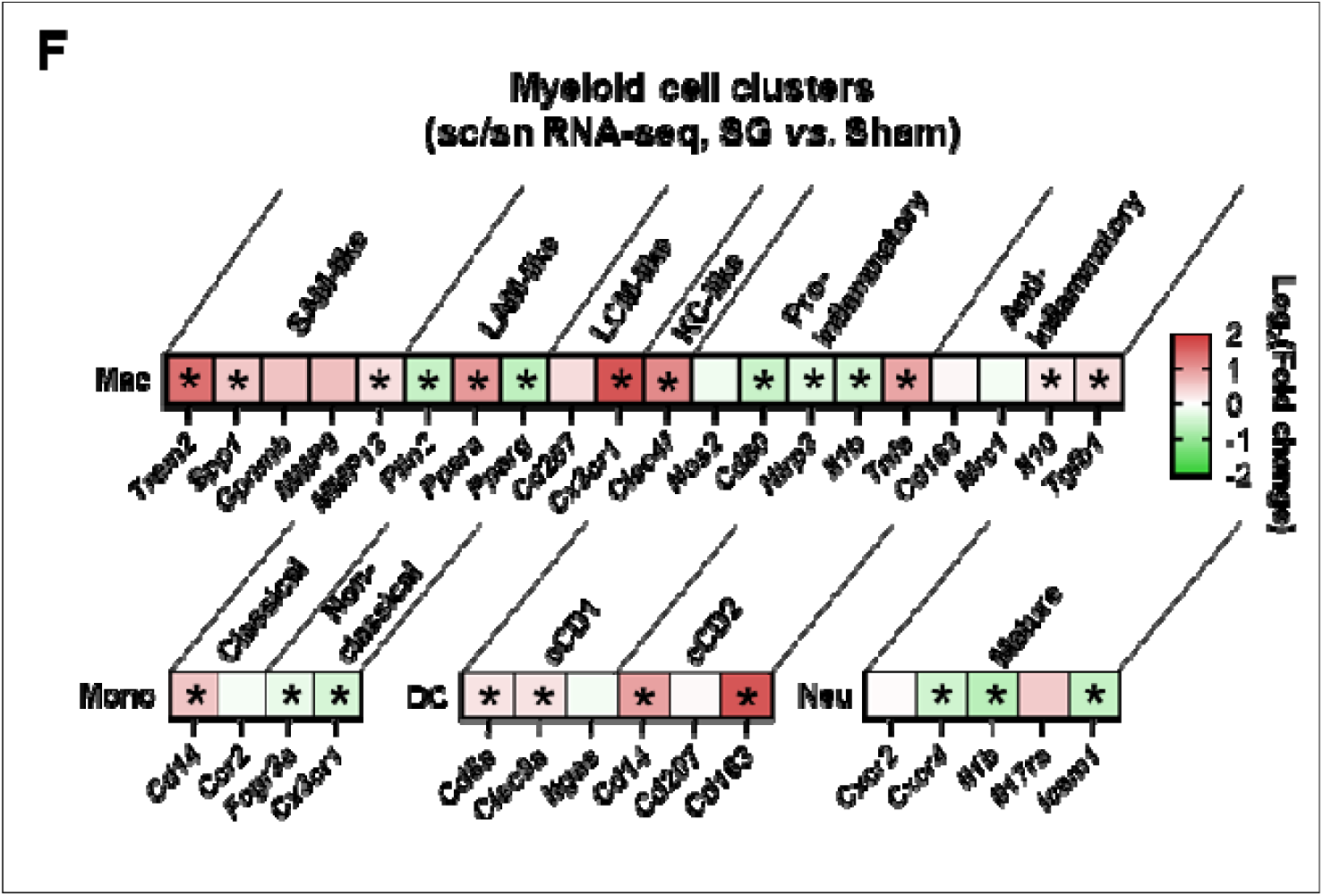
Sleeve gastrectomy facilitates lipid metabolism and phenotypic alterations in myeloid cells. **(A)** The technical scheme of the deep-learning-based image analysis. The deep-learning approach assessed **(B)** proportions of hepatocytes, lymphocytes and myeloid cells in total liver cells per tissue section and **(C)** spatial distribution in representative liver samples. Regions with higher (red circles) and lower (green circles) myeloid cell densities were selected **(D)** Differentially generated metabolites were illustrated (higher myeloid cell infiltrated regions *vs.* lower myeloid cell infiltrated regions) in Sham, Sham + DR and SG groups. **(E)** Function enrichment of upregulated differentially generated metabolites (higher myeloid cell infiltrated regions *vs.* lower myeloid cell infiltrated regions) in the liver sample of SG group was analyzed based on the KEGG database. **(F)** The expression of phenotype-associated markers in myeloid cell clusters were illustrated based on sc/sn RNA-seq data (SG *vs.* Sham). Abbreviations: C: control; Sham/S: sham surgery; DR: dietary restriction; SG: sleeve gastrectomy; KEGG: Kyoto encyclopedia of genes and genomes. sc/sn: single-cell/single-nuclei; Mono: monocytes; Mac: macrophages; DC: dendritic cells; Neu: neutrophils; SAM: scar-associated macrophage; LAM: lipid-associated macrophage; LCM: liver capsular macrophage; KC: Kupffer cell; cDC: conventional DC. The unpaired t-test and one-way ANOVA test were performed. ‘*’ represents ‘*p* < 0.05’ and statistical significance.

### SG facilitates liver repair processes in steatotic livers

The strong accumulation of macrophages in SG-treated livers, in conjunction with the weight development and metabolic improvement, suggested the induction of myeloid cell-driven hepatic repair processes ^22^. To further assess the cellular phenotypic alterations in the liver, a list of markers was explored (for hepatocytes: *Pcna*, *Ppara*, *Cyp2e1*, *Cyp7a1* and *Afp*; for cholangiocytes: *Pcna*, *Sox9*, *Slc4a2*, *Hnf4a* and *Hnf1b*; for HSCs: *Pdgfrb*, *Tgfb1*, *Col1a1*, *Acta2* and *Vim*) (Fig. 8A). The expression of *Pcna*, *Ppara*, *Cyp2e1* and *Cyp7a1* in hepatocytes, the expression of *Pcna* and *Hnf1b* in cholangiocytes, and the expression of *Pdgfrb* in HSCs were significantly upregulated by SG (*vs.* Sham). In contrast, the expression of *Pcna* in hepatocytes and cholangiocytes, together with the expression of *Pdgfrb* were downregulated by Sham + DR (*vs.* Sham) (Fig. 8A). Besides, cell differentiation was assessed and depicted using a cell trajectory analysis model (Fig. 8B). Cell trajectory of hepatocytes implied higher expression of *Pcna*, *Cyp2e1* and *Ppara* in Sham + DR and SG groups than the Sham group, while the expression distinctly varied with the cell trajectory (Fig. 8B, Fig. S3C). Cholangiocytes were classified into three groups. Cholangiocytes from Sham and SG groups were more associated with the expression of *Pcna* and *Slc4a2* but not with the expression of *Sox9* (Fig. 8B, Fig. S3C).

**Fig. 8.**
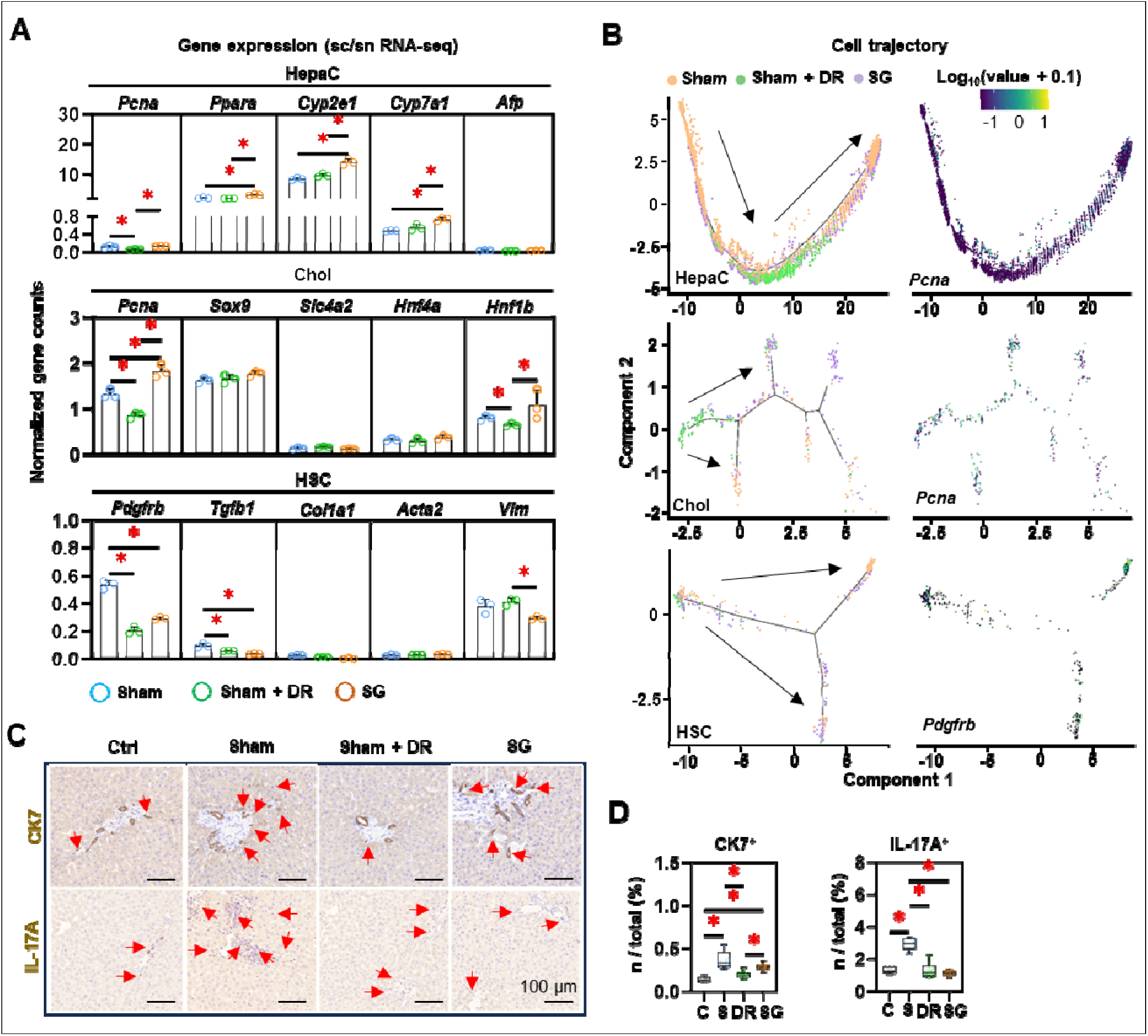
Dietary restriction and sleeve gastrectomy alter the cellular differentiation of hepatocytes, cholangiocytes and hepatic stellate cells. **(A)** Cellular characteristic markers of HepaC (*Pcna*, *Ppara*, *Cyp2e1*, *Cyp7a1* and *Afp*), Chol (*Pcna*, *Sox9*, *Slc4a2*, *Hnf4a* and *Hnf1b*) and HSC (*Pdgfrb*, *Tgfb1*, *Col1a1*, *Acta2* and *Vim*) were displayed. **(B)** Cell trajectories of HepaC, Chol and HSC together with *Cyp2e1* expression in HepaC, *Pcna* expression in Chol, and *Pdgfrb* expression in HSC. CK7- and IL-17A-expressing cells were **(C)** illustrated in situ and **(D)** quantified from total rat liver cells. Abbreviations: C: control; Sham/S: sham surgery; DR: dietary restriction; SG: sleeve gastrectomy; HepaC: hepatocyte; Chol: cholangiocyte; HSC: hepatic stellate cell; KEGG: Kyoto encyclopedia of genes and genomes. The one-way ANOVA test was performed. ‘*’ represents ‘*p* < 0.05’ and statistical significance.

Furthermore, HSCs of Sham + DR groups were shown in relatively inactivated clusters with low *Pdgfrb* expression (Fig. 8B), but not correlated with the expression of *Tgfb1* and *Col1a1* (Fig. S3B). In addition, CK7-expressing cells accumulated more in Sham and SG groups (*vs.* Ctrl and *vs.* Sham + DR) (Fig. 8C and D). Simultaneously, interleukin (IL)-17A-expressing cells increased in Sham groups (*vs.* Ctrl) but decreased in Sham + DR and SG groups (*vs.* Sham) (Fig. 8C and D). Taken together, SG can facilitate liver repair processes by enhancing the proliferation of hepatocytes and cholangiocytes, as well as fueling the fibrogenesis of HSCs.

### DR affects zonation characteristics of periportal hepatocytes

Given the remarkable cell trajectory with variable expression of phenotypic markers, differential characteristic alterations in hepatocyte subtypes were hypothesized. Hence, hepatocytes were further dissected into subpopulations based on sc/sn RNA-seq dataset (Fig. 9A). Metabolism-related function enrichment analysis revealed two distinct groups: metabolism one group (Met1, including cluster 6, 7 and 10) enriched metabolism processes of glycolysis, lipids and drugs, while metabolism two groups (Met2, including clusters 1, 2, 3, 4, 5, 8 and 9) enriched fatty acid biosynthesis (Fig. 9B). Interestingly, illustrations of the gene expression (pericentral hepatocyte markers: *Ctnnb1*, *Cnne1*, *Lrp5*, *Lrp6*, *Myc* and *Glul*; periportal hepatocyte markers: *Cyp2f2* and *Alb*) further identified that the Met1 group harbors more periportal hepatocytes, while the Met2 group harbors more pericentral hepatocytes (Fig. 9C and D). Furthermore, cell trajectory analysis indicated the explicit differentiation status of these two metabolism groups, which supports the zonal classification (Fig. 9E). In comparison to periportal (Met1) hepatocytes, pericentral (Met2) hepatocytes illustrated higher numbers and strength of interactions with immune cells, while myeloid cells, T lymphocytes and NK cells exerted particularly high involvement in cellular interactions (Fig. S8A and B). Particularly, periportal hepatocytes accounted for 15 - 20% of total hepatocytes in both Sham and SG groups but were almost absent (0.4%) in the Sham + DR group (Fig. 9F). Strikingly, proliferative cells (Ki67+) were more present in the liver tissues of Sham and SG groups in comparison to Ctrl groups. Significantly, SG enhanced cell proliferation among all groups (Fig. 9G). In addition, metabolism markers [cytochrome (CYP)2E1 and CYP7A1] and cell proliferation (Ki67) were investigated in situ. CYP2E1 and CYP7A1 were determined to be more enriched surrounding central veins in Ctrl, Sham and SG groups, but to be homogeneously distributed in Sham + DR groups (Fig. 9G and H). In addition, the spatial distribution of fatty acid, BA and sphingolipid metabolites appeared to be enriched around periportal areas in Sham and SG groups but rather widely spread in Sham + DR groups (Fig. 9I). Taken together, DR reshapes hepatocyte zonation features potentially by depriving periportal hepatocyte characteristics, which may compromise the capacities of metabolism and regeneration in steatotic livers in comparison to SG.

**Fig. 9.**
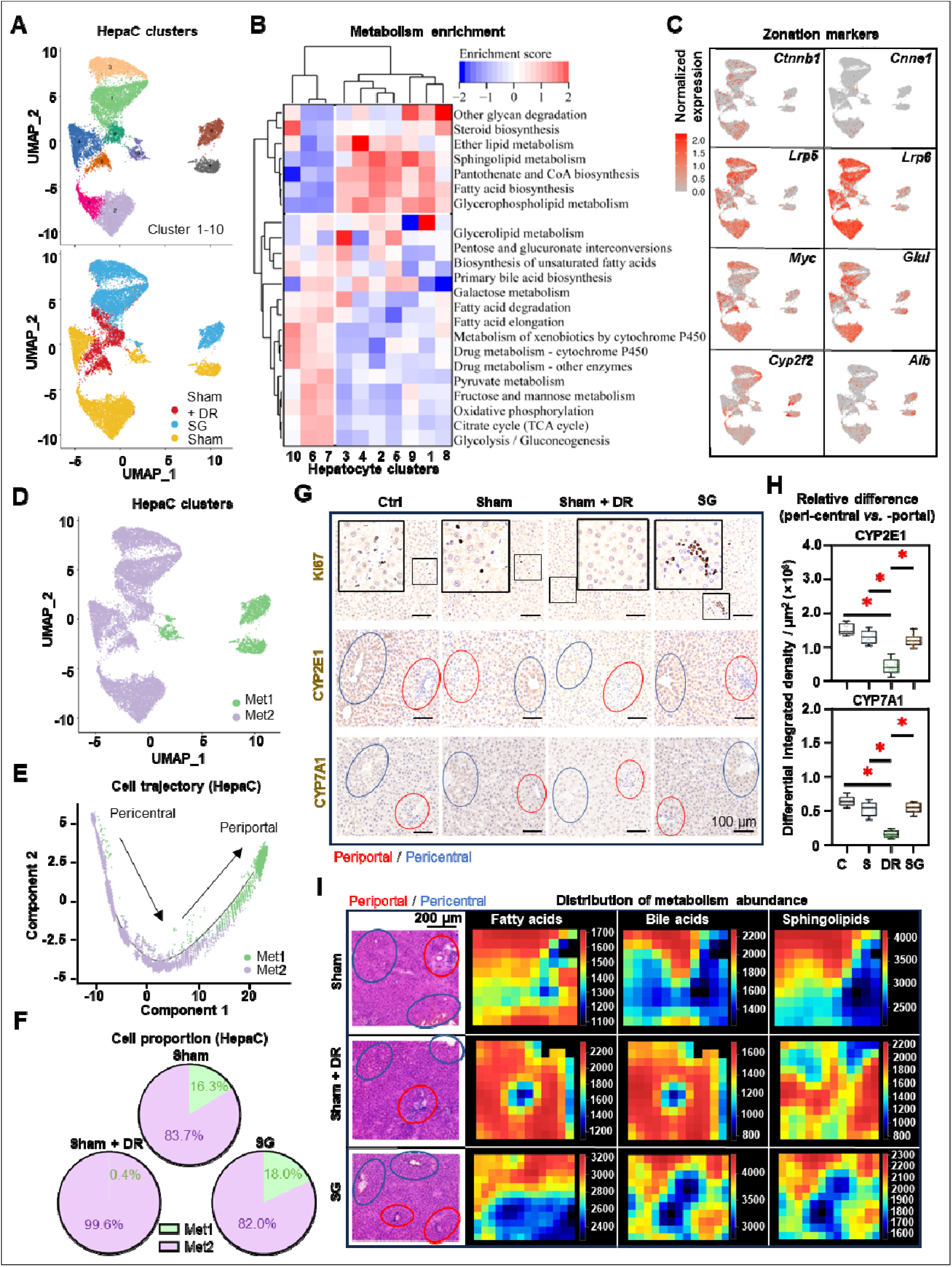
Dietary restriction compromises zonation characteristics of periportal hepatocytes. **(A)** Subclusters of rat hepatocytes from Sham, Sham + DR and SG illustrated in UMAPs. **(B)** Metabolism enrichment on each hepatocyte subcluster is displayed in a matrix. **(C)** Gene expression of *Ctnnb1*, *Cnne1*, *Lrp5*, *Lrp6*, *Myc*, *Glul*, *Cyp2f2* and *Cyp7a1* illustrated in rat hepatocytes. **(D)** Metabolism group-1 and −2 in rat hepatocytes. **(E)** Metabolism group-1 and −2 in the cell trajectory of rat hepatocytes. **(F)** Proportions of metabolism group-1 and −2 in rat hepatocytes of Sham, Sham + DR and SG groups. **(G)** Protein levels of Ki67, CYP2E1 and CYP7E1 illustrated in rat liver tissue (each view contains one or more portal and central areas) of Sham, Sham + DR and SG groups. **(H)** Relative differences of CYP2E1 and CYP7E1 levels between peri-central and -portal regions in each rat liver tissue of Sham, Sham + DR and SG groups. **(I)** Spatial distribution of fatty acid, bile acid and sphingolipid metabolites. In H&E- and IHC-characterized rat liver sample sections, periportal regions were outlined in red circles and pericentral regions were outlined in blue circles. Abbreviations: Sham: sham surgery; DR: dietary restriction; SG: sleeve gastrectomy; HepaC: hepatocyte; Met: metabolism; CYP: cytochrome. H&E: hematoxylin and eosin. ‘*’ represents ‘*p* < 0.05’ and statistical significance.

## Discussion

In this study, we employed multiomics methods to unravel relevant alterations of obesity-related parameters and liver steatosis following SG and DR in both patient cohorts and rat models. Specifically, sc/sn RNA-seq data unveiled an augmentation of lipid metabolism in both hepatocytes and immune cells, with a notable emphasis on myeloid cells following SG. Furthermore, spatial metabolomics analysis revealed the strengthened metabolic capacities of liver cells induced by SG and DR. Notably, SG elicited elevated lipid metabolic capacities in regions exhibiting intense myeloid cell infiltration, underscoring the intricate interplay between metabolic and immune responses in the liver microenvironment. PPAR-α emerged as a prominent candidate mediating the observed metabolic enhancements induced by both SG and DR. Additionally, the increased serological levels of bile acids and gut-derived FGF-19, together with hepatic butanoate metabolism underscored the pivotal roles of the gut-liver axis in modulating liver metabolism post-SG.

To date, only a few studies have employed the bulk transcriptomic analysis of patients’ liver samples post-BS to identify alterations of inflammation and lipid metabolism, suggesting potential therapeutic mechanisms of MASLD ^23^. Essentially, PPARα was elucidated as a key nuclear receptor that regulates weight-loss-induced MASLD amelioration ^24,25^. Consequently, PPAR agonism has been regarded as a promising approach to tackling MASLD ^26,27^. Robust preclinical data have determined the potential of targeting PPARα to improve lipid metabolism, inflammation, and fibrosis in MASLD ^28,29^. Our previous study has proved that the elevating PPAR levels by lanifibranor (a pan-PPAR agonist) significantly improves liver steatosis and fibrosis in mice and in an in-vitro liver-on-a-chip model of MASLD ^30^. However, numerous PPAR dual/pan agonists have failed to reach clinical practices, because they could not satisfy clinical endpoints (amelioration of e.g., liver injury, fibrosis, steatosis) ^31^. It has been reported that lipid metabolism is critical in immune response ^32,33^. Conspicuously, our study outlined plausible direct links between metabolism restoration and myeloid cell infiltration in post-SG steatotic rat livers. It has been known that myeloid cells, especially macrophages, play a pivotal role in pathological and therapeutic mechanisms of metabolic liver diseases ^27,34,35^. Liver macrophages can mediate hepatic lipid metabolism during steatotic stress ^36,37^, while lipid metabolism is closely associated with phenotypic alterations in macrophages ^38,39^. Thus, the liver macrophage remains a novel target for MASLD therapies ^40–42^. In addition, SG effectively enhanced the PPARα level and metabolic pathways broadly on hepatic parenchymal and immune cells, to a higher degree than DR. It has been demonstrated that PPARα activation in the liver can ameliorate the accumulation of circulating monocytes upon DR intervention ^43^. In addition, cell types in pathological rat liver samples were characterized using an AI-assisted digital method, which has been taken as an advanced tool for MASLD diagnosis ^18,44^. Intriguingly, in rat MASLD models, SG can strengthen the macrophage pool despite a broad PPARα activation in the liver. Further characterization of the macrophage population indicates that SG endows the anti-inflammatory and repair-associated potential of macrophages rather than detrimental pro-inflammatory roles in the liver.

The hepatic metabolic and regenerative functions vary by hepatocyte zonation (periportal and pericentral), leading to diverse responses to injuries and therapies ^45^. Emerging studies demonstrated that absent metabolic function in the periportal zone provokes cholestasis and ductular reaction ^46^. On the other hand, compromised hepatic periportal characteristics may be attributed to dysregulated WNT signaling and cell regeneration ^46,47^. Nowadays, the advances in spatially resolved multiomics approaches (e.g., sc/sn RNA-seq, spatial metabolomics, and spatial proteomics) provide both broader views and deeper understandings of heterogenic pathomechanisms of liver diseases ^45,48,49^. Our study elucidated that hepatic periportal characteristics were dramatically subsided by DR, whereas preserved by SG. Since DR is rarely associated with adverse events in the liver, we hypothesized that periportal hepatocytes might acquire ‘pericentral’ features to aid hepatic metabolism. In contrast, enhanced immunometabolism levels may interpret the preservation of zonal features upon the SG intervention. In addition, our data indicated that the regenerative potentials of hepatocytes were enhanced by SG but not DR, which strengthened the findings from the latest study ^17^.

Noteworthy, our study determined improved capacities of BA secretion and butanoate metabolism, as well as elevated FGF19 levels in serum upon the SG, suggesting that SG influences liver metabolism by regulating the gut-liver axis. Given the essential roles of gut-derived signals in the liver, the current understanding of MASLD pathogenesis lacks insights into gut-liver crosstalk, encompassing hepatic cell metabolism, inflammation, fibrogenesis, and complex cellular interactions, which likely play pivotal roles in MASLD progression ^22,27,50^. Increasing evidence indicates critical roles that BS may play in regulating the gut-liver axis, including GM, and BA metabolism ^51^. As suggested by recent studies, post-BS changes in the GM and BA circulation, as well as a decrease in the portal influx of free fatty acids, are beneficial to steatotic livers and obesity ^52–54^. Furthermore, BS can drastically repopulate GM and reverse the primary/secondary BA ratio in the intraluminal ileal, inducing improvement in metabolic syndrome in patients with MASLD ^55^. On the other hand, the butanoate production by GM was reported to be associated with bile acids and mediating liver metabolism ^56^. Notably, butanoate production can be enhanced after bariatric surgery ^57^.

Nonetheless, the limitations inherent in the current study should be acknowledged. The exclusion of patients with MASLD and obesity who solely underwent DR stemmed from challenges associated with dietary normalization to SG and inherent biases in dietary quantification. Furthermore, the sample size utilized for spatial metabolomics analysis was relatively modest, potentially constraining the generalizability of the findings. Despite these limitations, this study highlights a prominent enhancement in liver immunometabolism upon SG. Notably, multi-omics approaches offer a comprehensive exploration of the cellular and metabolic landscape within steatotic livers, marking a pioneering endeavor in this domain. Beyond traditional analyses, the incorporation of sc/sn RNA transcriptome and spatially resolved metabolite distribution analyses provides unprecedented insights into the immunometabolism dynamics of MASLD post-DR and - SG.

## Materials and Methods

### Human cohort

The clinical retrospective study was conducted on 18 patients who received SG in Changzhou Second People’s Hospital (from January 2023 to July 2023). This cohort comprised n = 8 male (44.4%) and n = 10 female (55.6%) individuals at a median age of 34 years (range: 29 - 40). We included patients that fulfilled the following criteria: 1) BMI > 32.5 kg/m^2^; 2) Age: 16 – 65; 3) clinically diagnosed with MASLD (based on cardiometabolic risk factors and presence of steatosis on liver imaging; 4) diagnosed with type 2 diabetes or sleep apnea syndrome; 5) without a history of other hepatic or genetic illness; 6) received sleeve gastrectomy, without postoperative complications. Clinical indexes (e.g., BMI, serological indexes) were evaluated on the day before SG and 6 months post-surgery. This clinical retrospective study was approved by the ethics committee of the Affiliated Changzhou Second People’s Hospital of Nanjing Medical University (ID: [2023]KY124-01).

### Animal models and surgical interventions

Male wild-type SPRAGUE DAWLEY® rats aged 8 weeks were obtained from Laboratory Animal Co., Ltd. (Changzhou, China). Rats were bred under specific-pathogen-free conditions and fed with HFD (D12492, Research Diets, Inc., USA) for 12 weeks. Five rats received sham surgery followed by uncontrolled HFD, 9 rats received sleeve gastrectomy followed by uncontrolled HFD, and 8 rats received sham surgery followed by restricted HFD feeding (comparable to dietary intake of post-SG rats) for 8 weeks. Procedures of sleeve gastrectomy were illustrated in supplementary data (Supplementary methods and Fig. S1) and were based on our previous research ^58^. Five healthy control rats were set with normal diet feeding but without any specific interventions. All these procedures were performed and supported by Kerbio Co., Ltd., Changzhou, China. Animal experiments were approved by the Institutional Animal Care and Use Committee (ID: IACUC23-0068).

### Serological tests

Patients’ blood samples were acquired from the median cubital vein. Rat blood samples were harvested through rat hearts immediately after rats were sacrificed. Serum was harvested using centrifugation (450×g, 10 min), and stored at −80LJ. Circulating markers including AST, ALT, PPG, γ-GT, TG, cholesterol (CHO), HDL-C, LDL-C, alkaline phosphatase (ALP) and TBA were measured using Indiko analyzer (ThermoFisher, USA). Cytokines including IL-6, IL-10, FGF-19, FGF-21 and transferrin (TRF) were measured using ELISA kits and a microplate reader (Rayto, China). All measurements were conducted following the manufacturer’s instructions. Antibodies that were used in this study are listed in Table S1.

### Immunohistochemistry and histological exploration

Liver tissues were harvested immediately after rats were sacrificed. Tissues were fixed in 10%-formalin for 24 hours, followed by paraffin embedding. Sections of 5 µm thickness were obtained using a HistoCore BIOCUT microtome (Leica, Germany). Hematoxylin and eosin (HE) staining were performed using standard procedures. Extracellular collagen area was indicated using the Picro Sirius Red Stain Kit (Abcam, UK). IHC method was employed using the kits purchased from Beyotime Biotechnology, China. Antigen retrieval was conducted using a heat-induced epitope retrieval method within a citrate buffer. Primary antibodies against the target proteins were applied, followed by appropriate secondary antibodies. Diaminobenzidine (DAB) was used for chromogenic detection, and counterstaining was performed with hematoxylin. Slides were examined under a microscope to evaluate protein localization and expression levels. In situ staining was observed under a light microscope (Olympus, Japan). Antibodies used in this study are listed in Table S1. For the quantitation of cell nucleus and protein levels, FIJI software was utilized ^59^. In particular, relative staining densities of proteins per µm2 were assessed and compared between peri-central and -portal regions in each tissue slide.

### Single-cell and single-nuclei sequencing and analysis

Single cells and cell nuclei were isolated and extracted from freshly collected rat liver samples using the dissection kit purchased from OE Biotech Co., Ltd., Shanghai, China. The libraries were acquired following the manufacturer’s protocol and then sequenced using the MobiNova platform (Shanghai, China). The integration of sc/sn-RNA sequencing data was operated following the common protocol with eliminating batch effects ^60^. Cell Ranger was used to filter the data, while the downstream analyses were conducted using Seurat (V5.0.1) ^61^, CellChat (1.6.0) ^62^, and Monocle2 ^63^ software packages for R. Function enrichment was performed based on Gene ontology (GO) and Kyoto Encyclopedia of Genes and Genomes (KEGG) databases. All the technical procedures including sampling, quality control, library build-up and sequencing analysis were supported by the genome platform of OE Biotech Co., Ltd., Shanghai, China.

### Spatial metabolomics analysis

Liver samples were freshly acquired from rats and then frozen at −80LJ immediately after optimal cutting temperature (OCT) embedding. Embedded tissue samples were sliced into 5 µm using a cryo-microtome CM 1950 (Leica, Germany). In-situ profiles of total metabolites were detected with positive and negative ion binding approaches using the MSI metabolomics method ^64^. Tissue regions were annotated after H&E staining. Metabolite enrichment was assessed based on the KEGG database. All the technical procedures including sample preparation, quality control, detection and analysis were supported by the metabolomics platform of Luming Biotechnology Co., Ltd. Shanghai, China.

### AI-guided imaging analysis

The AI-guided imaging analysis performed herein is based on a previously published approach ^65–67^. H&E-stained rat liver sections were scanned using a light microscope (Olympus, Japan). Processing of the whole slide images (WSIs) was operated using QuPath 0.4.4 following a pre-defined protocol ^68^. Sparse annotations, comprising approximately 15 cells for each of the cell populations, were manually labeled across three WSIs representing different conditions (Sham, Sham + DR and SG). A simple artificial neural network classifier with two hidden layers, consisting of 20 and 10 neurons, respectively, was trained to distinguish between the three classes versus the rest. Example regions without these classes were used as negative examples during training. Subsequently, cell classification and measurements were exported, and for each cell population, 2D hexagonal binning plots of the number of cells were generated in Python (using the matplotlib function ‘hexbin’) ^69^. The model was trained for representative cells (at least 15 cells/sample’ of hepatocytes, lymphocyte-like cells and myeloid-like cells) based on three liver tissue slides, which were defined with the assistance of pathological expertise. The trained model was applied to the sample cohort of rat liver sections (including 5 Ctrl, 5 Sham, 8 Sham + DR and 8 SG samples). Detailed methodology is described in the Supplementary methods.

### Statistical analysis

The GraphPad Prism 9.0 software (GraphPad Software, USA) and the ggplot2 package (version: 3.4.4) for R were used to generate plots ^70^. The statistical tests used in this study are indicated in the figure legend for each panel. Data are presented as the mean ± S.D. ‘*p* < 0.05’ was considered a significant difference.

## Acknowledgments

Xiurong Cai and Hanyang Liu were supported by China Scholarship Council Foundation. The authors thank Shanghai OE Biotech CO., Ltd. and Shanghai Luming Biological Technology Co., Ltd. for the access to the MobiNova single-cell system and AFADESI-MSI platforms. Schematic figures were generated via BioRender.

## Funding

National Nature Science Foundation of China (82300730)

Changzhou Health Commission of China (CZQM2022007 and QN202121)

Changzhou Science and Technology Bureau of China (CJ20220142)

Changzhou Medical Center of Nanjing Medical University (CZKYCMCB202221)

German Research Foundation (DFG Ta434/8-1, SFB/TRR 296 and CRC1382, Project-ID 403224013).

## Author contributions

H.L. conceptualized the research. H.L. and A.G. supervised the research. S.C., Q.Z. and H.L. developed methodology. S.C. and P.S. performed animal experiments. Q.Z. and C.K. performed computational analysis. S.C. and J.X. performed experiments and clinical data collection. H.L., X.C. and S.C. performed data analysis. The manuscript was written by H.L. and revised by A.G. and F.T. H.L., A.G. and F.T. acquired funding.

## Data and materials availability

Raw and processed data of sc/sn-RNA sequencing analysis can be obtained from the GEO database (https://www.ncbi.nlm.nih.gov/geo/) with the reference number GSE254841. Raw data of spatial metabolomics can be obtained from the OMIX, China National Center for Bioinformation (https://ngdc.cncb.ac.cn/omix/) with the reference number OMIX005793. Technical materials for the deep-learning-based image analysis model may be provided by the corresponding authors upon reasonable requests. All data are available in the main text or the supplementary materials.

