## Supplementary Table 1 for "Multiomics analyses decipher intricate changes in the cellular and metabolic landscape of steatotic livers upon dietary restriction and sleeve gastrectomy"

**Table S1. Clinical follow-up data** (One day pre-SG vs. six months post-SG)

| Total | | | |
|---|---|---|---|
| **Variables** | **pre-SG (N = 18)** | **post-SG (N = 18）** | ***p*-value** |
| **BMI** | 37.01 (34.96 - 39.05) | 28.70 (26.81 - 30.58), ↓ | <0.05 |
| **PPG** (mmol/L) | 9.23 (13.44 - 8.01) | 7.23 (9.21 - 7.05), ↓ | <0.05 |
| **AST** (IU/L) | 38.80 (26.64 - 50.96) | 16.56 (13.45 - 19.67), ↓ | <0.05 |
| **ALT** (IU/L) | 67.28 (38.41 - 96.15) | 16.44 (9.03 - 23.85), ↓ | <0.05 |
| **γ-GT** (IU/L) | 57.89 (34.84 - 80.94) | 15.89 (11.49 - 20.28), ↓ | <0.05 |
| **TG** (mmol/L) | 1.84 (1.25 - 2.44) | 1.02 (0.88 - 1.17), ↓ | <0.05 |
| **CHO** (mmol/L) | 4.82 (4.33 - 5.32) | 4.95 (4.49 - 5.41), ↑ | 0.400 |
| **HDL-C** (mmol/L) | 1.19 (1.05 - 1.33) | 1.35 (1.21 - 1.48), ↑ | <0.05 |
| **LDL-C** (mmol/L) | 2.96 (2.49 - 3.43) | 3.13 (2.67 - 3.59), ↑ | 0.419 |
| **TBIL** (μmol/L) | 10.67 (8.51 - 12.84) | 14.19 (10.20 - 18.18), ↑ | <0.05 |
| **ALP** (IU/L) | 72.83 (61.04 - 84.63) | 74.00 (65.62 - 82.38), ↑ | 0.849 |
| **TBA** (μmol/L) | 2.84 (1.53 - 4.16) | 4.88 (0.75 - 10.51), ↑ | 0.051 |
| **TRF** (g/L) | 41.01 (35.21 - 46.81) | 36.20 (31.78 - 40.62), ↓ | 0.175 |
| **IL-6** (pg/mL) | 398.18 (316.07 - 480.29) | 485.43 (385.89 - 584.96), ↑ | 0.072 |
| **IL-10** (pg/mL) | 33.98 (32.18 - 35.79) | 36.55 (34.24 - 38.86), ↑ | 0.626 |
| **FGF-19** (pg/mL) | 491.23 (178.42 - 768.31) | 642.16 (255.67 - 880.72), ↑ | 0.058 |
| **FGF-21** (pg/mL) | 1219.67 (1210.03 - 1349.31) | 1326.99 (1247.35 - 1419.62), ↑ | 0.378 |

Continued

| | Male | | |
|---|---|---|---|
| **Variable** | **pre-SG (N = 8)** | **post-SG (N = 8)** | ***p*-value** |
| **BMI** | 39.4 (37.4 - 41.0) | 30.8 (29.8 - 32.4), ↓ | <0.05 |
| **PPG** (mmol/L) | 9.31 (13.44 – 8.24) | 7.78 (9.21 – 7.15), ↓ | <0.05 |
| **AST** (IU/L) | 34 (30 - 63) | 16 (14 - 18), ↓ | <0.05 |
| **ALT** (IU/L) | 55 (48 - 122) | 16 (13 - 18), ↓ | <0.05 |
| **γ-GT** (IU/L) | 82 (35 - 107) | 24 (14 - 27), ↓ | <0.05 |
| **TG** (mmol/L) | 2.01 (1.52 - 3.61) | 1.05 (0.93 - 1.21), ↓ | <0.05 |
| **CHO** (mmol/L) | 4.90 (4.55 - 4.99) | 5.32 (4.98 - 5.74), ↑ | 0.059 |
| **HDL-C** (mmol/L) | 1.05 (0.85 - 1.20) | 1.22 (1.06 - 1.31), ↑ | 0.301 |
| **LDL-C** (mmol/L) | 2.86 (2.54 - 3.12) | 3.71 (3.52 - 4.04), ↑ | <0.05 |
| **TBIL** (μmol/L) | 12 (9 - 14) | 15 (10 - 21), ↑ | 0.403 |
| **ALP** (IU/L) | 74 (58 - 78) | 78 (63 - 84), ↑ | 0.414 |
| **TBA** (μmol/L) | 3 (2 - 3) | 1 (1 - 3), ↓ | 0.103 |
| **TRF** (g/L) | 40 (36 - 47) | 36 (29 - 41), ↓ | 0.210 |
| **IL-6** (pg/mL) | 434 (316 - 456) | 423 (342 - 577), ↓ | 0.703 |
| **IL-10** (pg/mL) | 34.2 (32.9 - 35.79) | 35.5 (34.24 – 37.79), ↑ | 0.654 |
| **FGF-19** (pg/mL) | 465.43 (178.42 - 723.42) | 631.22 (255.67 - 856.23), ↑ | 0.234 |
| **FGF-21** (pg/mL) | 120 (120 - 129) | 132 (126 - 136), ↑ | 0.304 |

Continued

| Female | | | |
|---|---|---|---|
| **Variable** | **pre-SG (N = 10)** | **post-SG (N = 10)** | ***p*-value** |
| **BMI** | 35.1 (32.0 - 36.7) | 26.9 (24.0 - 30.2), ↓ | <0.05 |
| **PPG** (mmol/L) | 9.17 (13.00 – 8.01) | 7.18 (8.89 – 7.05), ↓ | <0.05 |
| **AST** (IU/L) | 28 (17 - 41) | 15 (14 - 17), ↓ | <0.05 |
| **ALT** (IU/L) | 39 (22 - 63) | 13 (8 - 15), ↓ | <0.05 |
| **γ-GT** (IU/L) | 32 (20 - 42) | 10 (8 - 10), ↓ | <0.05 |
| **TG** (mmol/L) | 1.10 (0.95 - 1.45) | 0.93 (0.77 - 1.03), ↓ | 0.121 |
| **CHO** (mmol/L) | 4.86 (4.10 - 5.26) | 4.70 (3.85 - 5.30), ↓ | 0.903 |
| **HDL-C** (mmol/L) | 1.33 (1.18 - 1.41) | 1.41 (1.27 - 1.50), ↓ | 0.210 |
| **LDL-C** (mmol/L) | 2.93 (2.48 - 3.62) | 3.09 (2.15 - 3.34), ↑ | 0.721 |
| **TBIL** (μmol/L) | 9.55 (6.90 - 12.40) | 11.25 (8.98 - 13.57), ↑ | 0.304 |
| **ALP** (IU/L) | 78 (60 - 93) | 77 (66 - 80), ↓ | 0.700 |
| **TBA** (μmol/L) | 1.90 (1.45 - 2.28) | 2.25 (1.30 - 2.88), ↑ | 0.701 |
| **TRF** (g/L) | 34 (29 - 49) | 32 (30 - 38), ↓ | 0.913 |
| **IL-6** (pg/mL) | 307 (253 - 583) | 448 (362 - 552), ↑ | 0.314 |
| **IL-10** (pg/mL) | 33.6 (32.18 - 34.21) | 36.7 (35.2 - 38.86), ↑ | >0.9 |
| **FGF-19** (pg/mL) | 520.13 (213.10 - 768.31) | 672.98 (270.81 - 880.72), ↑ | 0.050 |
| **FGF-21** (pg/mL) | 1235 (1149.03 – 1336.03) | 1311 (1204.10 – 1509.03), ↑ | 0.509 |

Values are presented as 'mean (minimum - maximum)'.
**Abbreviation**: SG: sleeve gastrectomy; BMI: body mass index; PPG: postprandial blood glucose; AST: aspartate aminotransferase; ALT: alanine aminotransferase; γ-GT: gamma-glutamyltransferase; TG, triglycerides; CHO, cholesterol; HDL-C, high-density lipoprotein cholesterol; LDL-C, low-density lipoprotein cholesterol; TBIL, total bilirubin; ALP, alkaline phosphatase; TBA, total bile acid. TRF: transferrin; IL: interleukin; FGF, fibroblast growth factor. The paired t-test was used. ‘$p < 0.05$’ is considered to be statistically significant.
