## Supplementary Table 2 for "Multiomics analyses decipher intricate changes in the cellular and metabolic landscape of steatotic livers upon dietary restriction and sleeve gastrectomy"

**Table S2. List of antibodies used in this study.**

| Method | Antigen | Manufacturer | Catalog No. | Clone | Host species | Dilution |
|---|---|---|---|---|---|---|
| IHC | PPAR-α | ThermoFisher | 600-401-421 | Polyclonal | Rabbit | 1/200 |
| | PNPLA3 | ThermoFisher | 67369-1-IG | 1D2B11 | Mouse | 1/200 |
| | FXR | ThermoFisher | 417200 | A9033A | Mouse | 1/200 |
| | CK7 | ThermoFisher | MA1-06315 | RCK105 | Mouse | 1/500 |
| | IL-17A | ThermoFisher | PA5-79470 | Polyclonal | Rabbit | 1/200 |
| | Ki67 | ThermoFisher | MA5-14520 | SP6 | Rabbit | 1/500 |
| | CYP2E1 | ThermoFisher | PA5-52652 | Polyclonal | Rabbit | 1/500 |
| | CYP7A1 | ThermoFisher | PA5-100892 | Polyclonal | Rabbit | 1/200 |
| ELISA | IL-6 | ThermoFisher | BS-0782R | Polyclonal | Rabbit | 1/1000 |
| | IL-6 | ThermoFisher | M620 | 5IL6 | Mouse | 1/1000 |
| | IL-10 | ThermoFisher | PA5-95561 | Polyclonal | Rabbit | 1/1000 |
| | TRF | ThermoFisher | A1-46375 | 57-6 | Mouse | 1/1000 |
| | FGF-19 | ThermoFisher | PA5-79252 | Polyclonal | Rabbit | 1/1000 |
| | FGF-19 | Meibiao biology | MB-7355B | Polyclonal | Rabbit | 1/1000 |
| | FGF-21 | ThermoFisher | PA5-79255 | Polyclonal | Rabbit | 1/1000 |

**Abbreviation**: PPAR: peroxisome proliferator-activated receptor; PNPLA3: patatin-like phospholipase domain containing protein 3; FXR: farnesoid X receptor; CK7: cytokeratin 7; IL: interleukin; FGF: fibroblast growth factor; CYP: cytochrome P450; TRF: transferrin.
