## Supplementary methods and figures for "Multiomics analyses decipher intricate changes in the cellular and metabolic landscape of steatotic livers upon dietary restriction and sleeve gastrectomy"

### SUPPLEMENTARY MATERIALS

#### Supplementary methods

1. Animal modeling and surgery procedures

SPRAGUE DAWLEY® male rats, aged 8 weeks, were purchased from Cavens Laboratory Animal Co., Ltd. After one week of adaptation feeding, they were fed with a high-fat diet (HFD; D12492, Research Diets, Inc., USA) for 12 weeks. They were randomly divided into three treatment groups: sleeve gastrectomy (SG), sham surgery, and sham surgery + diet restriction (DR). Before the surgery, the rats were fasted and deprived of water for 12 hours. They were then anesthetized with 1% pentobarbital sodium (4 mL/kg) intraperitoneal injection, placed on a fixed board, and the surgical area was disinfected. An incision of about 2 cm was made in the middle of the abdomen, and the stomach was located and pulled out from the abdominal cavity. Subsequently, the hepatic and splenic ligaments were ligated and sectioned, and the short gastric and gastrosplenic vessels were ligated. A pair of vascular forceps was applied from the cardia HIS Angle along the greater curvature of the stomach. The gastric body was quickly cut off with tissue scissors, and the incision of the gastric cavity was sutured continuously with 5 - 0 number sterile thread. The vascular clamp was then used to achieve control over the stomach, extending from the incision up to 3 mm above the pylorus. Approximately 70% to 80% of the gastric tissue, including the fundus, was excised, and subsequently, the incision was closed through continuous suturing. Finally, a varus suture was performed to minimize the risk of postoperative wound infection. The peritoneum, muscle, and skin were meticulously closed in layers to complete the sleeve gastric operation. Intraperitoneal injection of Meloxicam at a dosage of 0.5 mg/kg was administered for pain management on the first day after surgery (sham and SG). Additionally, rats preemptively received Penicillin at a dosage of 30,000 U/kg on the first day after surgery (sham and SG). Following the surgery, all rats underwent a one-day fasting period before being provided with sugar and saline on the second day post-operation. Subsequently, their diet gradually turned back to planned feeding.

The anesthesia and perioperative treatment of the sham group and sham + DR group were the same as that of the SG group. An incision of 1 cm was simply made in the stomach and sutured in place for the sham group. Additionally, the diet of the sham + DR group matched that of the SG group. The daily food intake of the SG group was measured, and the average amount was fed to the sham + DR group on the following day. The whole procedure is described in Figure S1.

2. AI-guided imaging analysis

The whole slide images (WSIs) with a magnification of 20x (0.2749 μm/pixel) were processed using QuPath 0.4.4 to measure hepatocyte, lymphocyte, and myeloid cell populations. Tissue detection was conducted by applying a threshold of 200 on the average of the red, green, and blue channels at a resolution of 12.5 μm/pixel on the smoothed image, followed by median filtering and smoothed coordinates. Tissue regions detected smaller than 1,000 px and holes smaller than 500 px were subsequently removed.

Nuclei were identified based on the hematoxylin optical density at a resolution of 1.5 μm/pixel, using a background radius of 8 and a sigma of 1.5. Following the opening morphological operation, objects detected smaller than 10 $\mu m^2$ or larger than 500 $\mu m^2$ were excluded from further analysis. The intensity threshold was set to 0.1, with a maximum background intensity of 2. Additionally, the boundaries of the detected nuclei were smoothed.

Sparse annotations, comprising approximately 15 cells for each of the cell populations, were manually labeled across three WSIs representing different conditions (Sham, DR and SG). A simple artificial neural network classifier with two hidden layers, consisting of 20 and 10 neurons, respectively, was trained to distinguish between the three classes versus the rest. Example regions without these classes were used as negative examples during training. Subsequently, cell classification and measurements were exported, and for each cell population, 2D hexagonal binning plots of the number of cells were generated in Python (using matplotlib function 'hexbin') with a blue-yellow-red color map for visualization purposes. The reported metrics included total cell counts of each cell type, as well as the ratio of each cell type to both the total tissue area and the total number of detected cells.

An analogous workflow was applied to 27 test WSIs. Considering slight color differences, the tissue detector utilized a threshold of 240 at a resolution of 20 µm/pixel, removing tissue regions smaller than 5,000 $\mu m^2$ and holes smaller than 1000 $px^2$. The same cell detector, cell classifier, and post-analysis procedures were employed for consistency.

**Supplementary figures and legends**

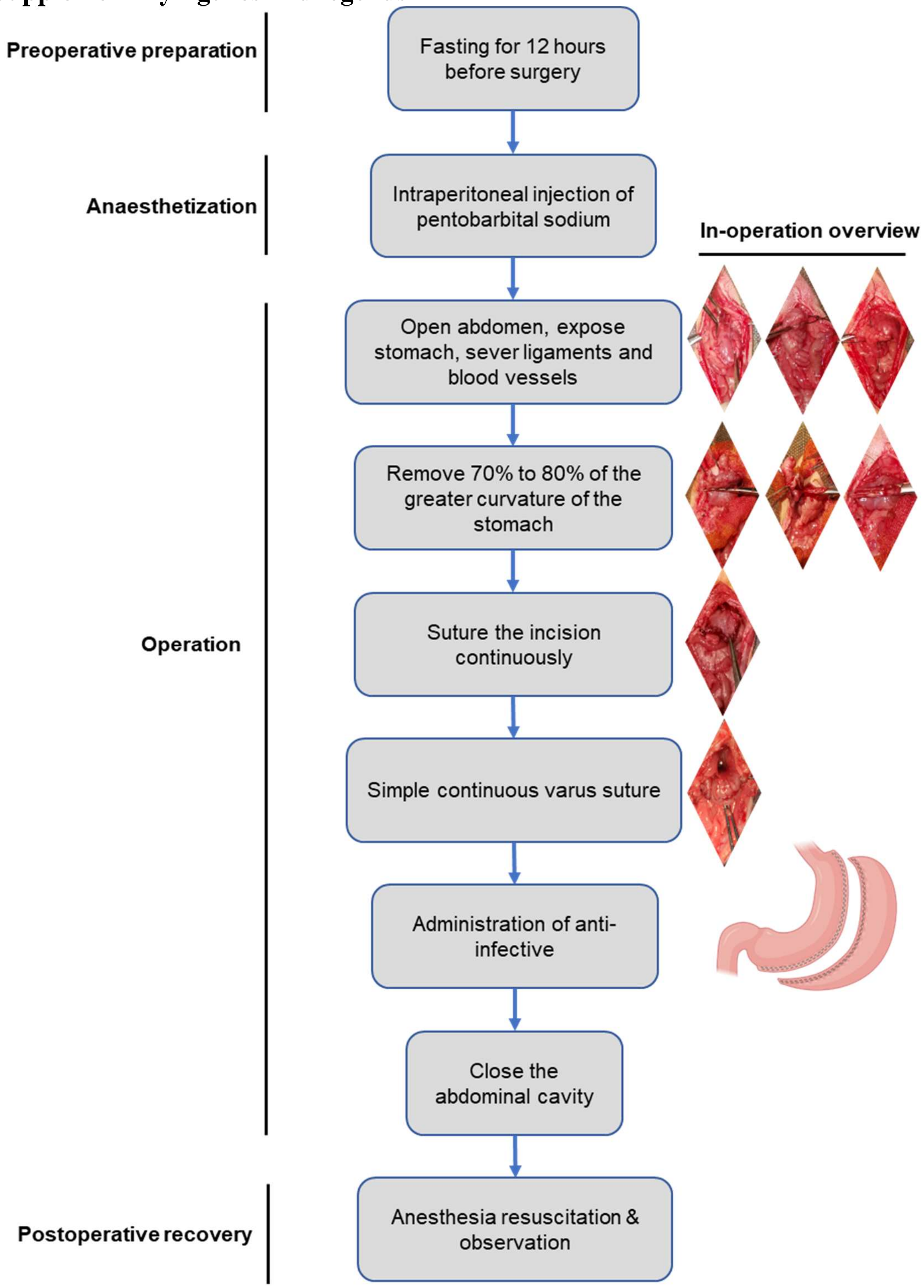

**Figure S1. Technical procedures of sleeve gastrectomy on rats.**

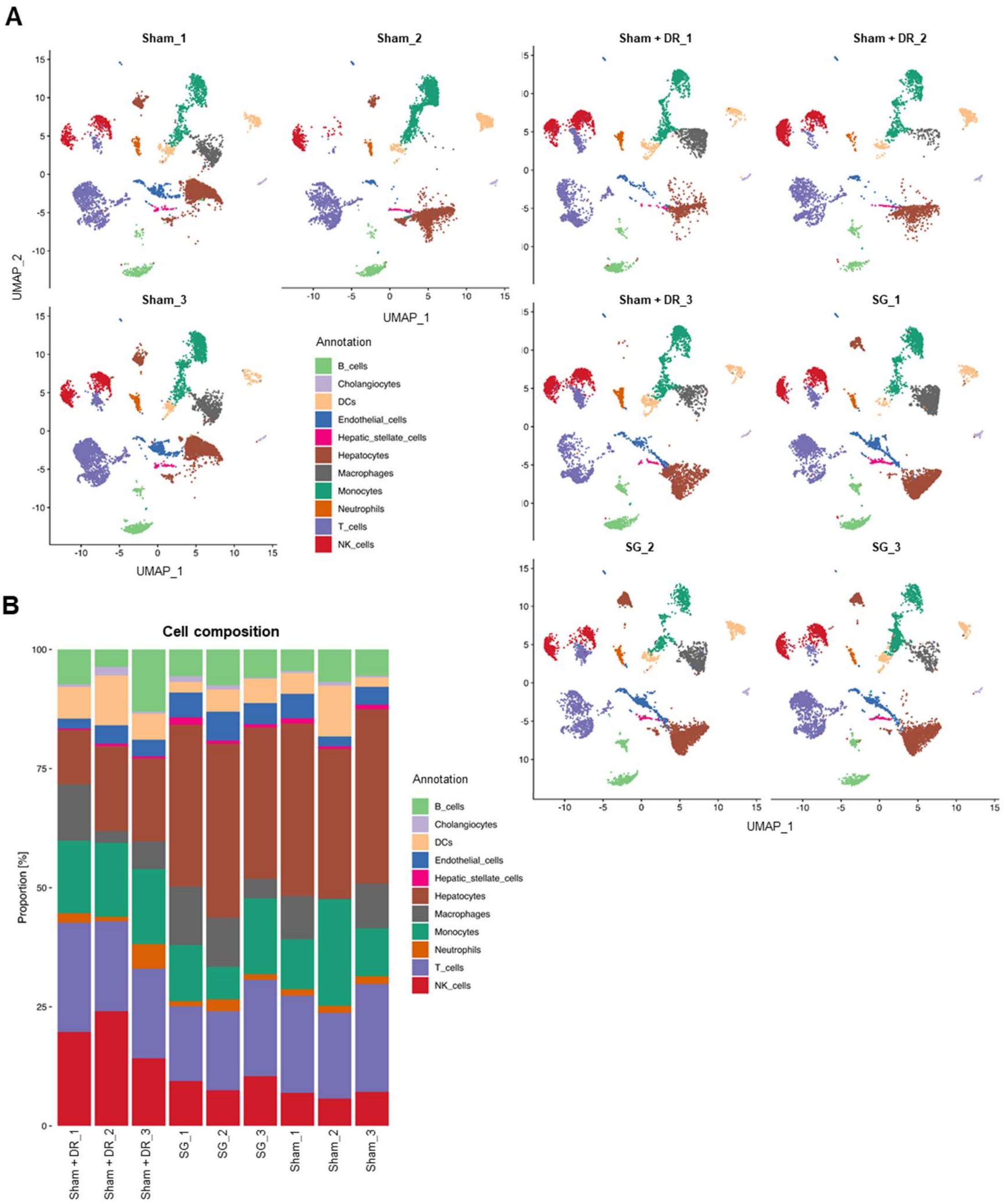

**Figure S2. Sc/sn RNA-seq analysis identifies cell populations. (A)** UMAPs illustrating the cell distribution of all individual samples upon Sham, Sham + DR and SG. **(B)** Histograms illustrating cell composition of all individual samples upon Sham, Sham + DR and SG. Abbreviations: Sham: sham surgery; DR: dietary restriction; SG: sleeve gastrectomy; B cells: B lymphocytes; T_cells: T lymphocytes; DC: dendritic cell.

A

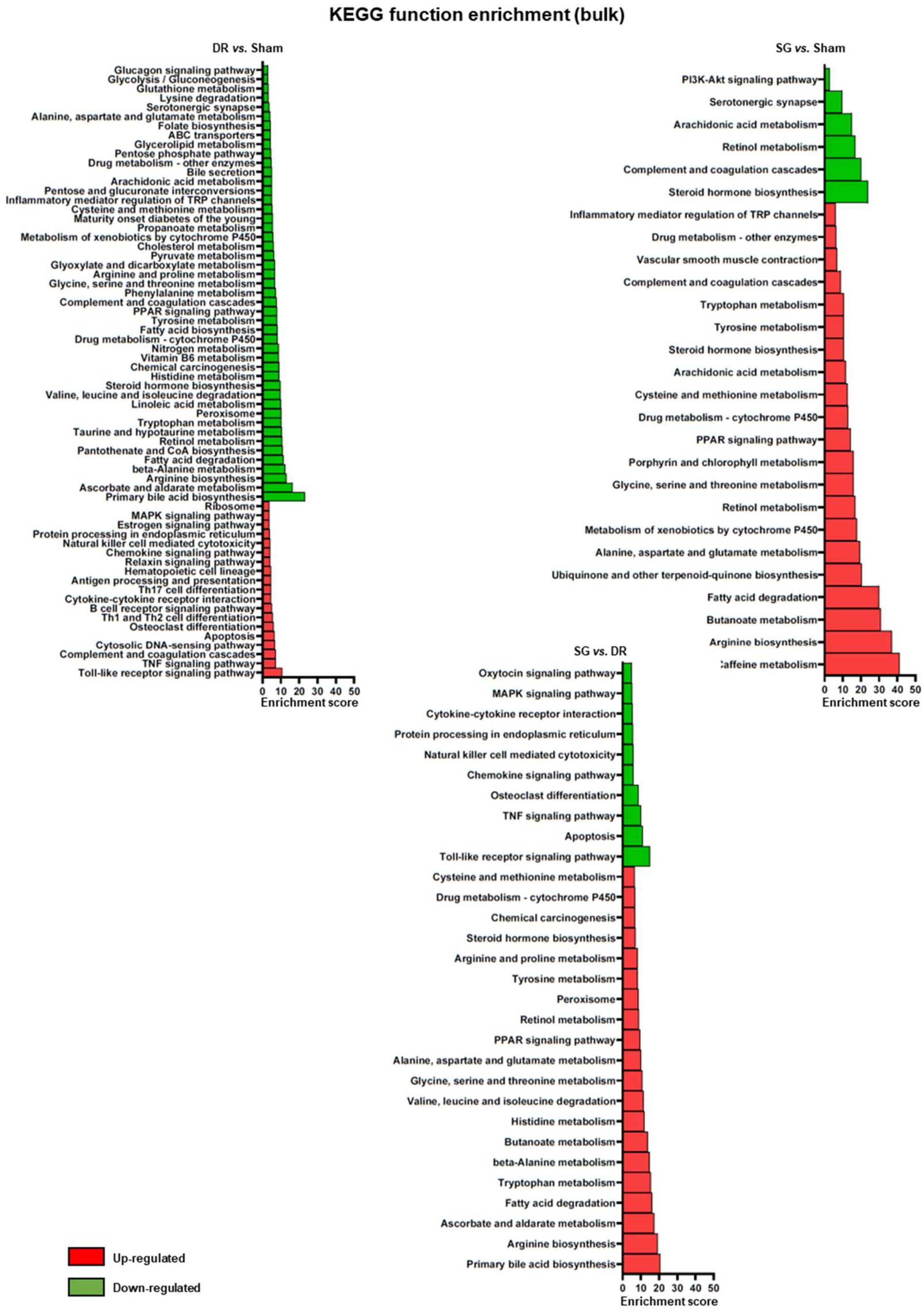
KEGG function enrichment (bulk)
DR vs. Sham
Glucagon signaling pathway
Glycolysis / Gluconeogenesis
Glutathione metabolism
Lysine degradation
Serotonergic synapse
Alanine, aspartate and glutamate metabolism
Folate biosynthesis
ABC transporters
Glycerolipid metabolism
Pentose phosphate pathway
Drug metabolism - other enzymes
Bile secretion
Arachidonic acid metabolism
Pentose and glucuronate interconversions
Inflammatory mediator regulation of TRP channels
Cysteine and methionine metabolism
Maturity onset diabetes of the young
Propanoate metabolism
Metabolism of xenobiotics by cytochrome P450
Cholesterol metabolism
Pyruvate metabolism
Glyoxylate and dicarboxylate metabolism
Arginine and proline metabolism
Glycine, serine and threonine metabolism
Phenylalanine metabolism
Complement and coagulation cascades
PPAR signaling pathway
Tyrosine metabolism
Fatty acid biosynthesis
Drug metabolism - cytochrome P450
Nitrogen metabolism
Vitamin B6 metabolism
Chemical carcinogenesis
Histidine metabolism
Steroid hormone biosynthesis
Valine, leucine and isoleucine degradation
Linoleic acid metabolism
Peroxisome
Tryptophan metabolism
Taurine and hypotaurine metabolism
Retinol metabolism
Pantothenate and CoA biosynthesis
Fatty acid degradation
beta-Alanine metabolism
Arginine biosynthesis
Ascorbate and aldarate metabolism
Primary bile acid biosynthesis
Ribosome
MAPK signaling pathway
Estrogen signaling pathway
Protein processing in endoplasmic reticulum
Natural killer cell mediated cytotoxicity
Chemokine signaling pathway
Relaxin signaling pathway
Hematopoietic cell lineage
Antigen processing and presentation
Th17 cell differentiation
Cytokine-cytokine receptor interaction
B cell receptor signaling pathway
Th1 and Th2 cell differentiation
Osteoclast differentiation
Apoptosis
Cytosolic DNA-sensing pathway
Complement and coagulation cascades
TNF signaling pathway
Toll-like receptor signaling pathway
0 10 20 30 40 50
Enrichment score
SG vs. Sham
PI3K-Akt signaling pathway
Serotonergic synapse
Arachidonic acid metabolism
Retinol metabolism
Complement and coagulation cascades
Steroid hormone biosynthesis
Inflammatory mediator regulation of TRP channels
Drug metabolism - other enzymes
Vascular smooth muscle contraction
Complement and coagulation cascades
Tryptophan metabolism
Tyrosine metabolism
Steroid hormone biosynthesis
Arachidonic acid metabolism
Cysteine and methionine metabolism
Drug metabolism - cytochrome P450
PPAR signaling pathway
Porphyrin and chlorophyll metabolism
Glycine, serine and threonine metabolism
Retinol metabolism
Metabolism of xenobiotics by cytochrome P450
Alanine, aspartate and glutamate metabolism
Ubiquinone and other terpenoid-quinone biosynthesis
Fatty acid degradation
Butanoate metabolism
Arginine biosynthesis
:affeine metabolism
0 10 20 30 40 50
Enrichment score
SG vs. DR
Oxytocin signaling pathway
MAPK signaling pathway
Cytokine-cytokine receptor interaction
Protein processing in endoplasmic reticulum
Natural killer cell mediated cytotoxicity
Chemokine signaling pathway
Osteoclast differentiation
TNF signaling pathway
Apoptosis
Toll-like receptor signaling pathway
Cysteine and methionine metabolism
Drug metabolism - cytochrome P450
Chemical carcinogenesis
Steroid hormone biosynthesis
Arginine and proline metabolism
Tyrosine metabolism
Peroxisome
Retinol metabolism
PPAR signaling pathway
Alanine, aspartate and glutamate metabolism
Glycine, serine and threonine metabolism
Valine, leucine and isoleucine degradation
Histidine metabolism
Butanoate metabolism
beta-Alanine metabolism
Tryptophan metabolism
Fatty acid degradation
Ascorbate and aldarate metabolism
Arginine biosynthesis
Primary bile acid biosynthesis
0 10 20 30 40 50
Enrichment score
Up-regulated
Down-regulated

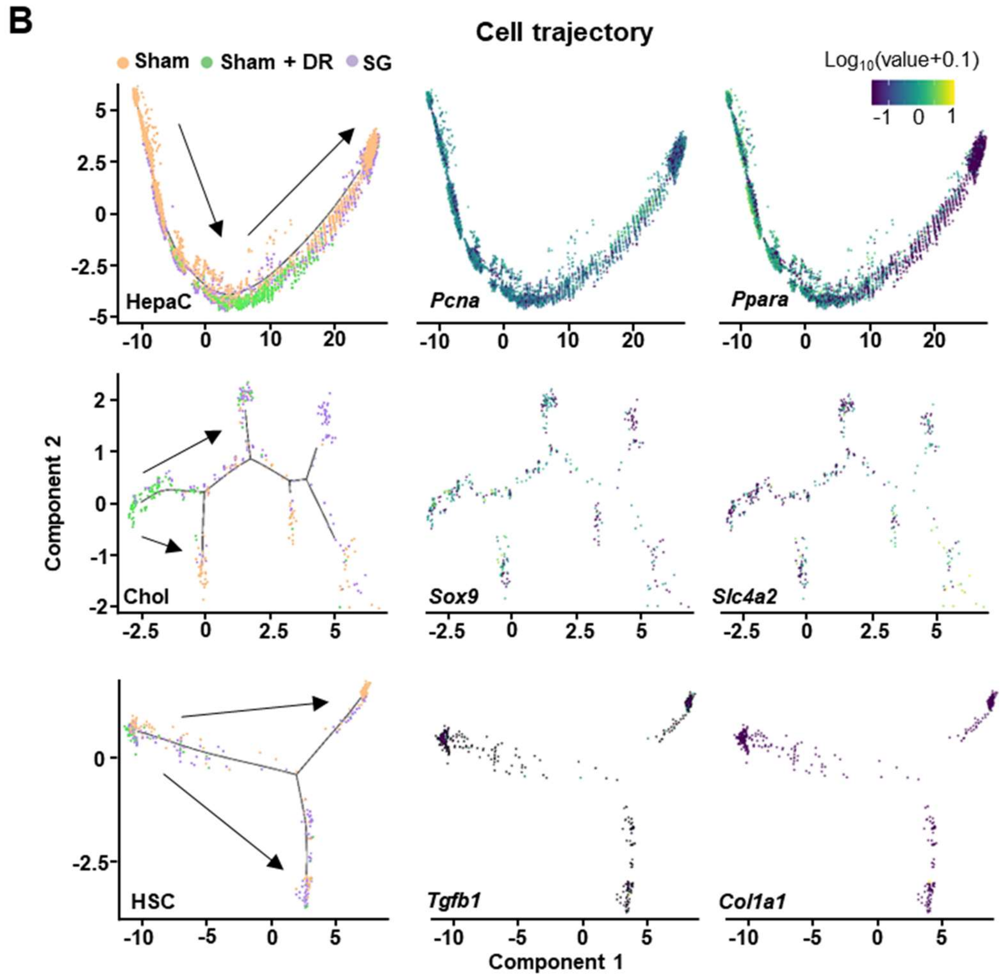

**Figure S3. Sc/sn RNA-seq analysis deciphers differentially expressed genes and cell trajectories**. **(A)** KEGG function enrichment analysis on significant DEGs upon comparisons (Sham + DR *vs.* Sham and SG *vs.* Sham). **(B)** Cell trajectories of HepaC, Chol and HSC, the gene expression of *Pcna* and *Ppara* in HepaC, the gene expression of *Sox9* and *Slc4a2* in Chol, and the gene expression of *Tgfb1* and *Col1a1* in HSC. Abbreviations: Sham: sham surgery; DR: dietary restriction; SG: sleeve gastrectomy; HepaC: hepatocytes; Chol: cholangiocyte; HSC: hepatic stellate cells; KEGG: Kyoto Encyclopedia of Genes and Genomes; DEG: differentially expressed gene.

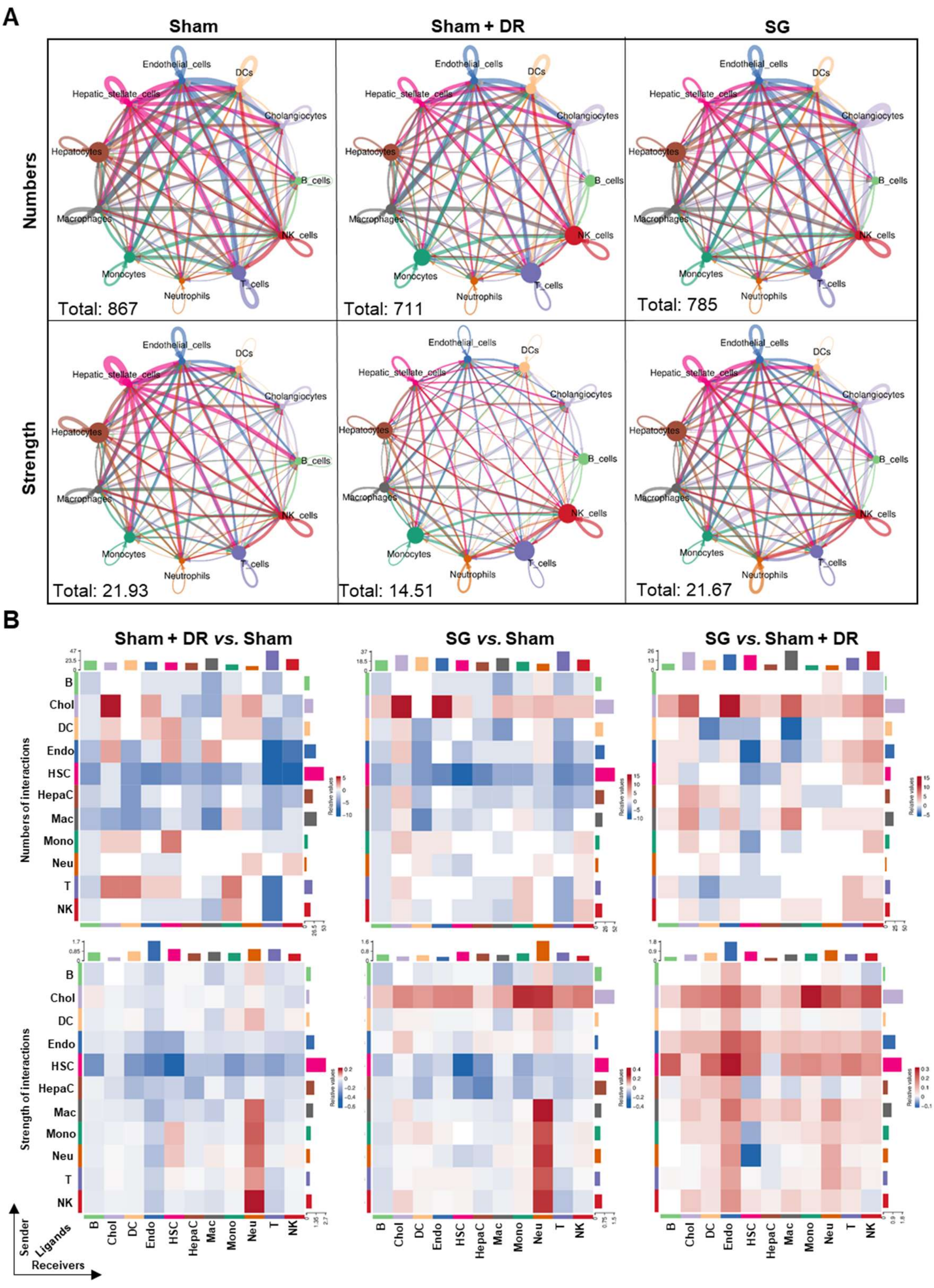

**Figure S4. Sc/sn RNA-seq analysis deciphers cellular interactions.** Numbers and strength of cellular interactions among cholangiocytes, endothelial cells, hepatic stellate cells, hepatocytes, B, T, NK, DCs, macrophages, monocytes and neutrophils in Sham, Sham + DR and SG groups illustrated in **(A)** networks and **(B)** matrixes. Abbreviations: Sham: sham surgery; DR: dietary restriction; SG: sleeve gastrectomy; DC: dendritic cells; NK: natural killer cells; B and B_cells: B lymphocytes; T and T_cells: T lymphocytes; Neu: neutrophils; Mac: macrophages; Mono: monocytes:

Endo: endothelial cells; HSC: hepatic stellate cells; Chol: cholangiocytes; HepaC: hepatocytes.

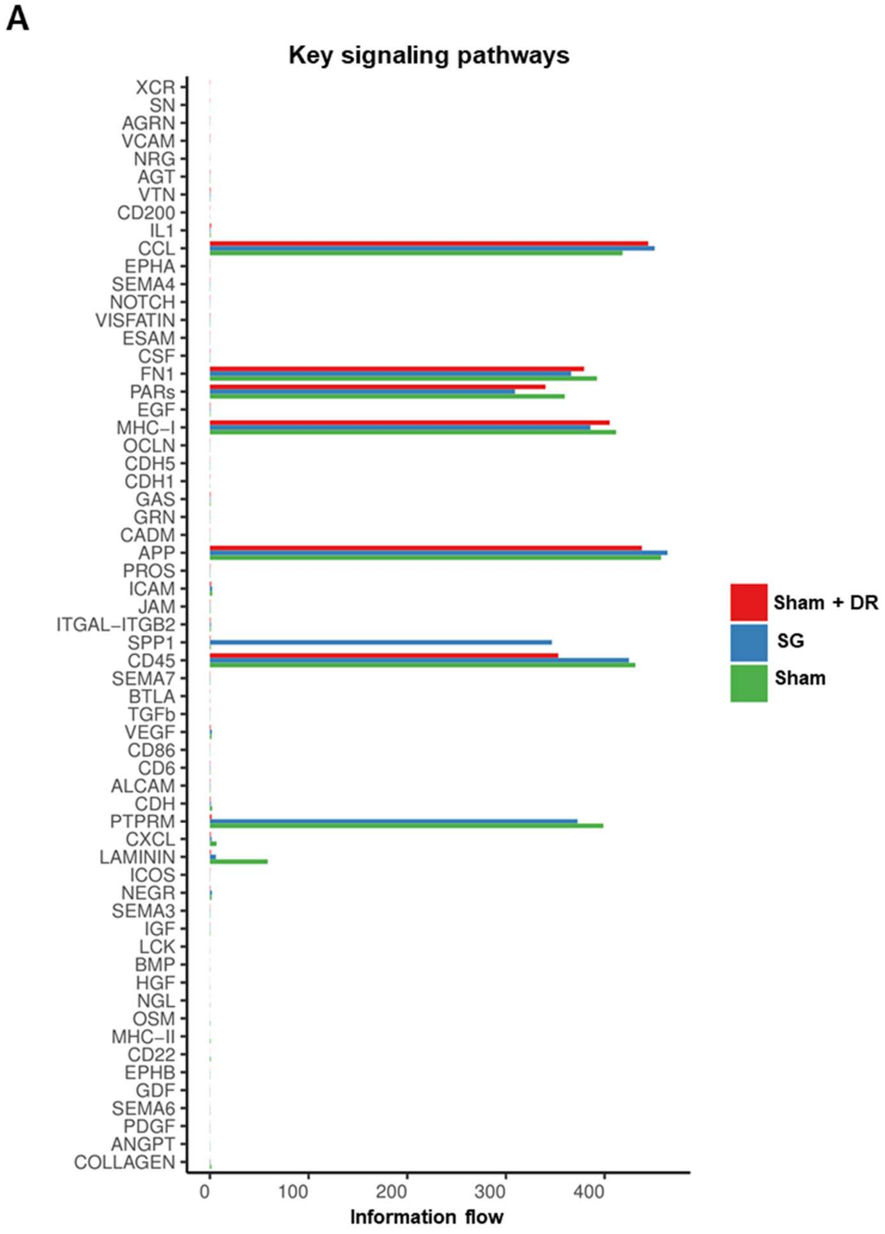

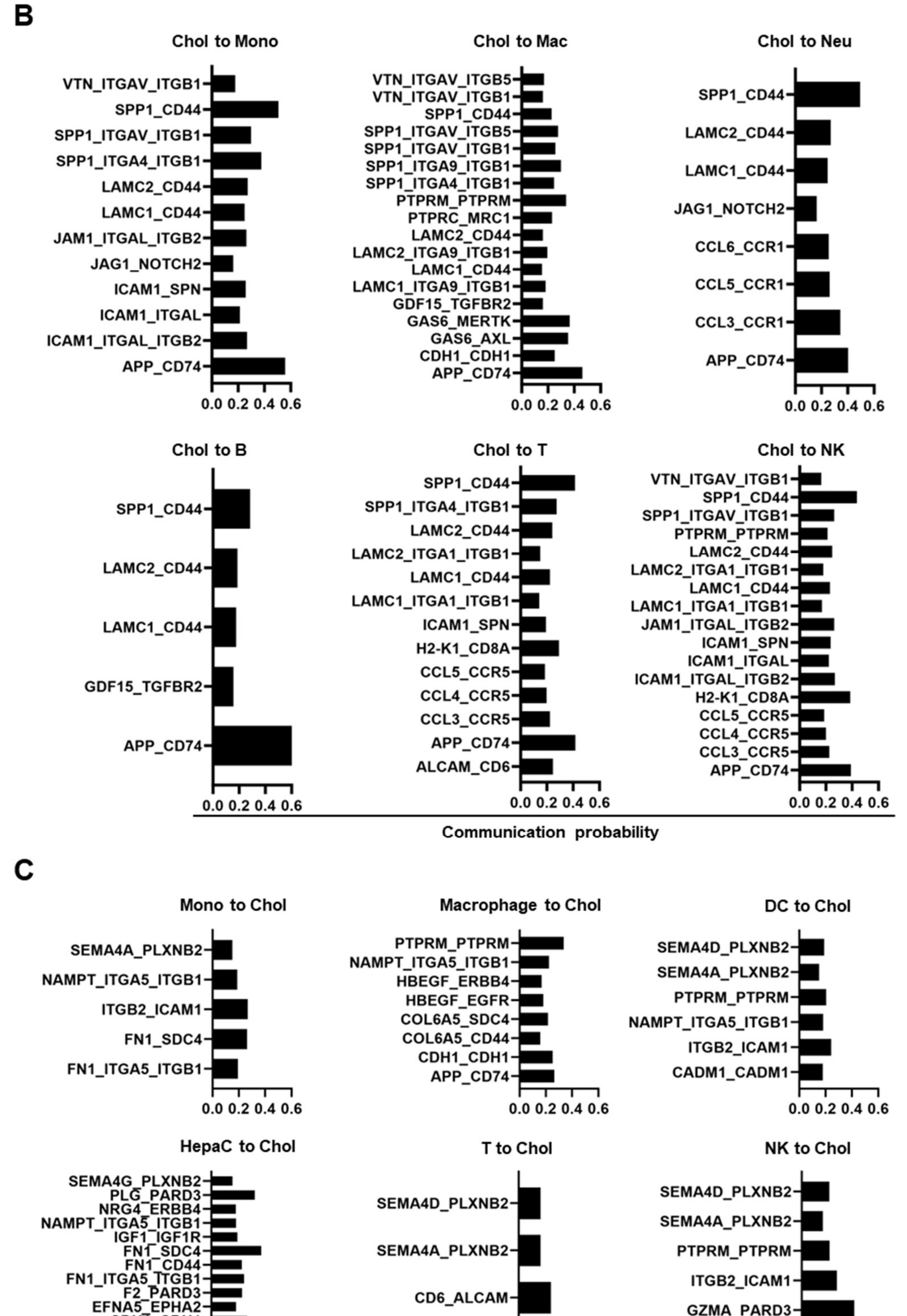

**Figure S5. Dissecting key pathways in cellular interactions. (A)** Frequency of key signaling pathways in Sham, Sham + DR and SG groups. Significant signaling pathways **(B)** from cholangiocytes to monocytes, macrophages, neutrophils, B, T and NK cells, as well as (C) from monocytes, macrophages, DCs, HepaCs, T and NK cells to cholangiocytes. Abbreviations: Sham: sham surgery; DR: dietary restriction; SG: sleeve gastrectomy; DC: dendritic cells; NK: natural killer cells; B: B lymphocytes; T:

T lymphocyte; Neu: neutrophils; Mac: macrophages; Mono: monocytes; Chol: cholangiocytes; HepaC: hepatocytes.

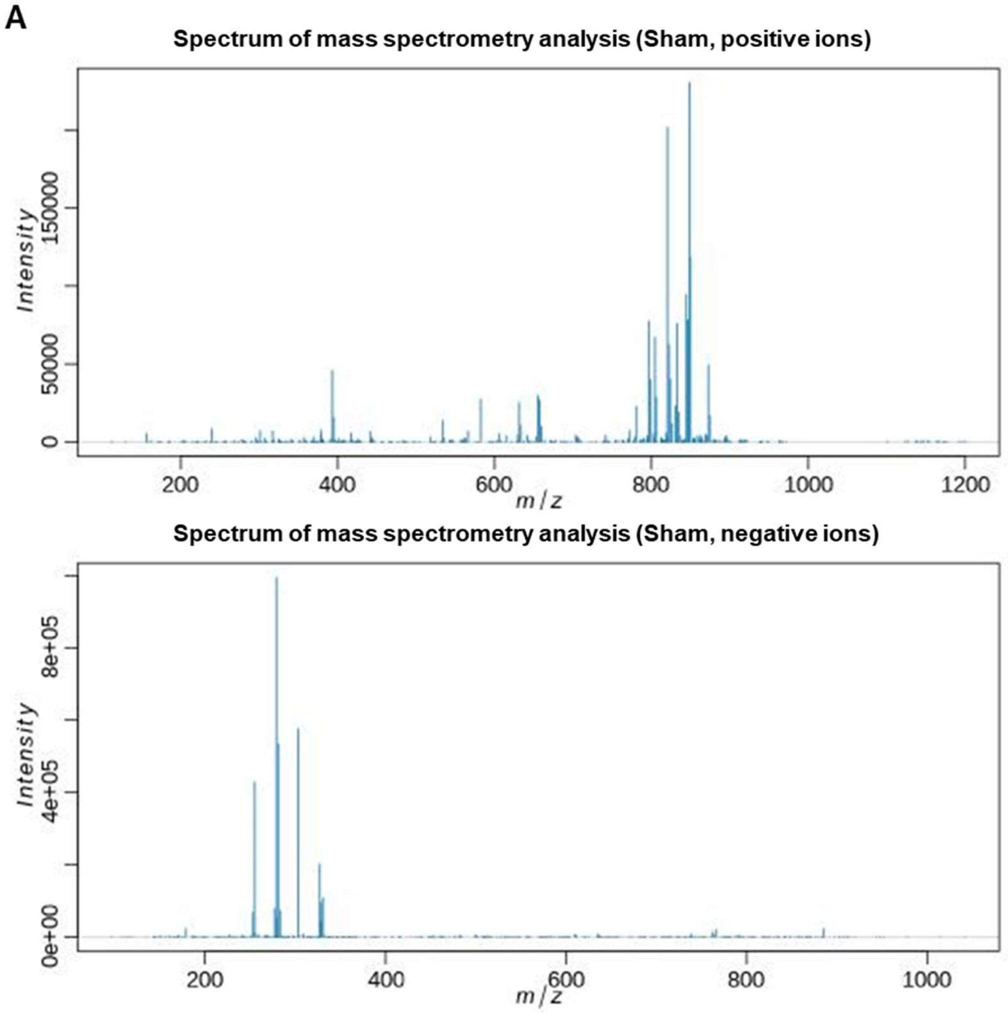

B

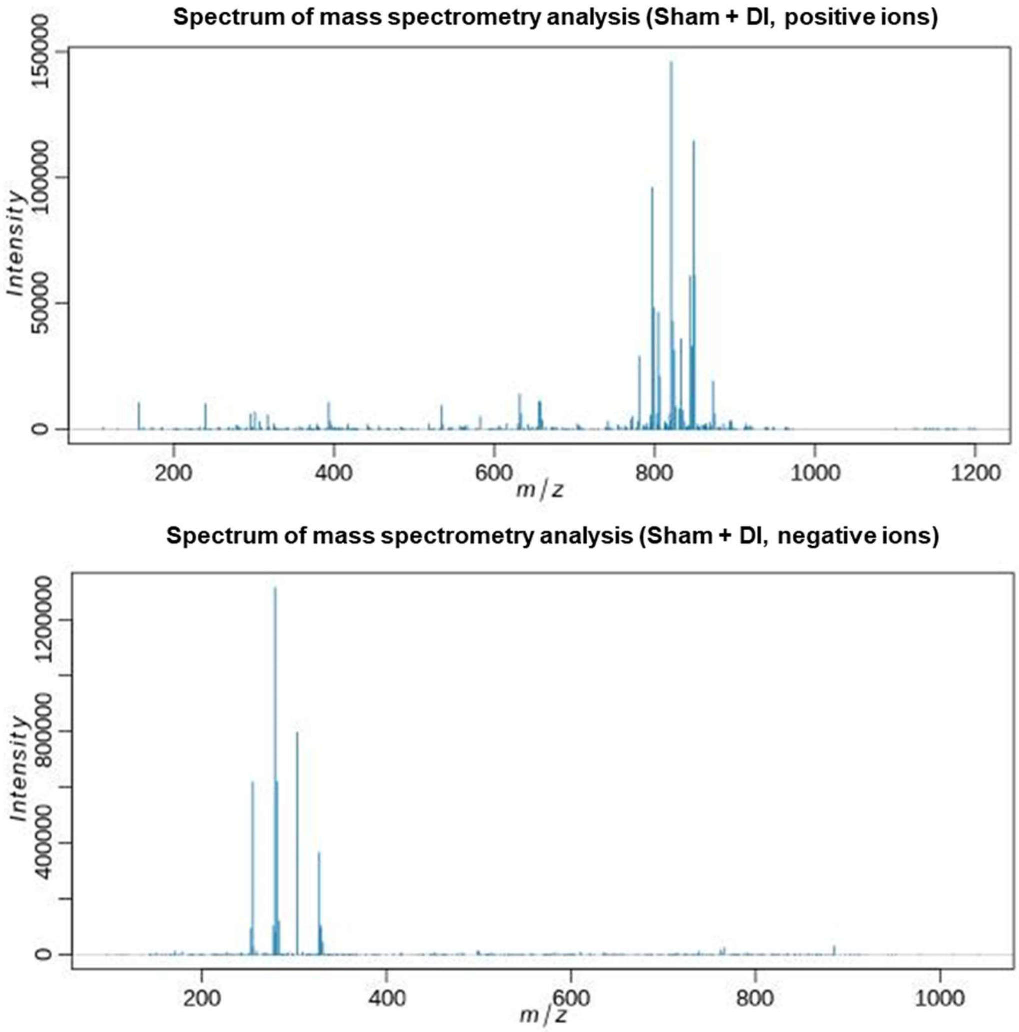

Spectrum of mass spectrometry analysis (Sham + DI, positive ions)
Intensity
0
50000
100000
150000
200
400
600
800
1000
1200
m/z
Spectrum of mass spectrometry analysis (Sham + DI, negative ions)
Intensity
0
400000
800000
1200000
200
400
600
800
1000
m/z

C

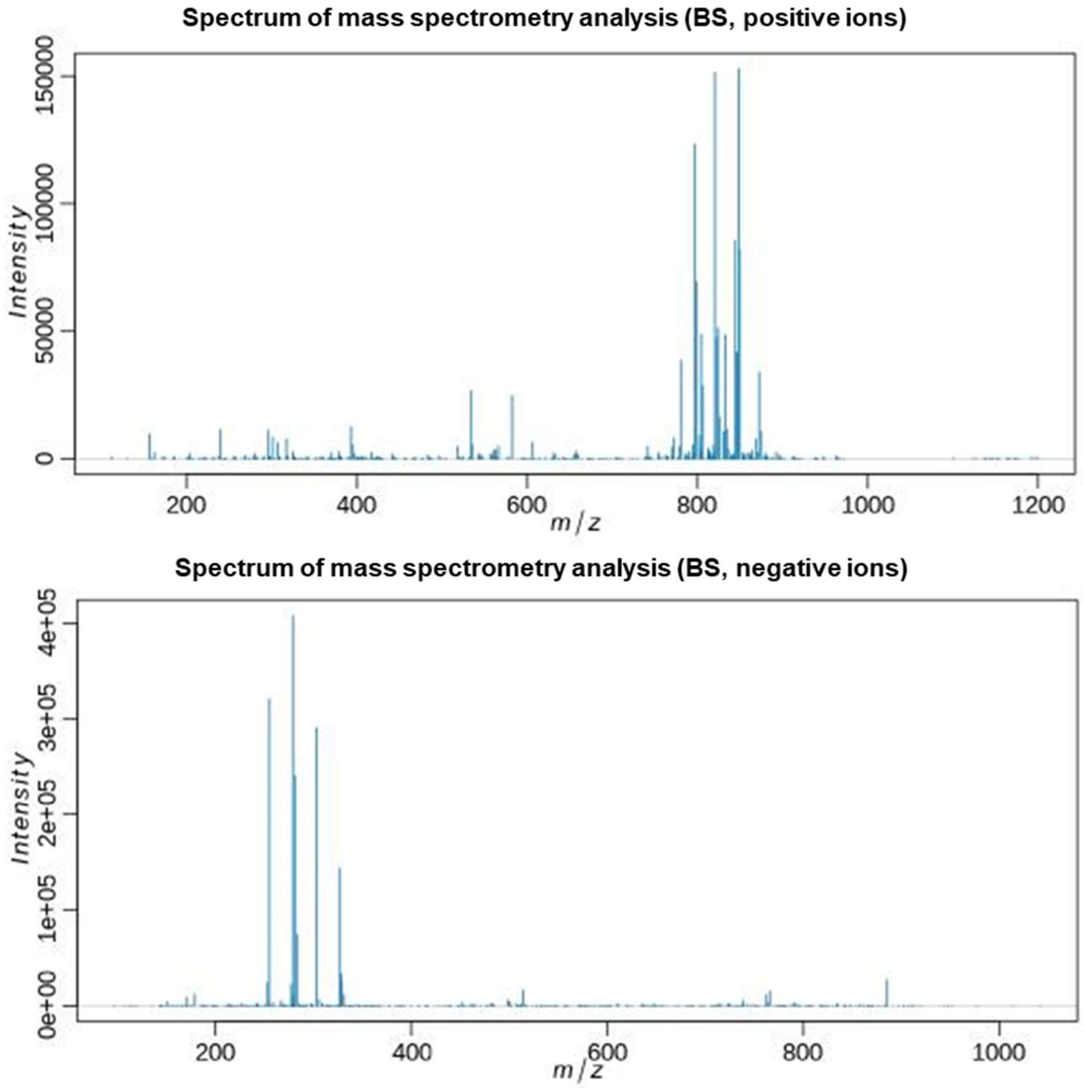
Spectrum of mass spectrometry analysis (BS, positive ions)
Intensity
0
50000
100000
150000
200
400
600
800
1000
1200
m/z
Spectrum of mass spectrometry analysis (BS, negative ions)
Intensity
0e+00
1e+05
2e+05
3e+05
4e+05
200
400
600
800
1000
m/z

**D**

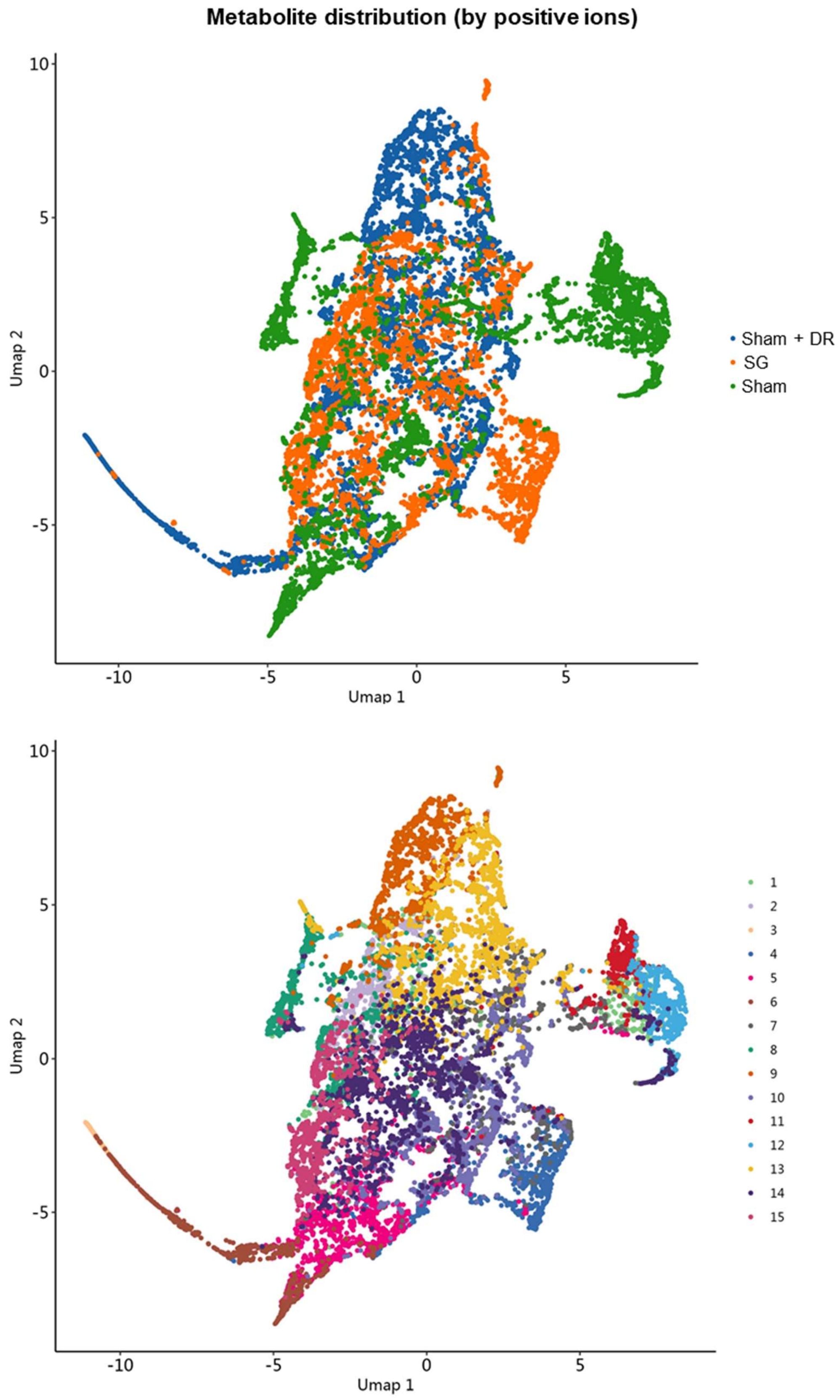
Metabolite distribution (by positive ions)
Umap 2
Umap 1
10
5
0
-5
-10
Sham + DR
SG
Sham

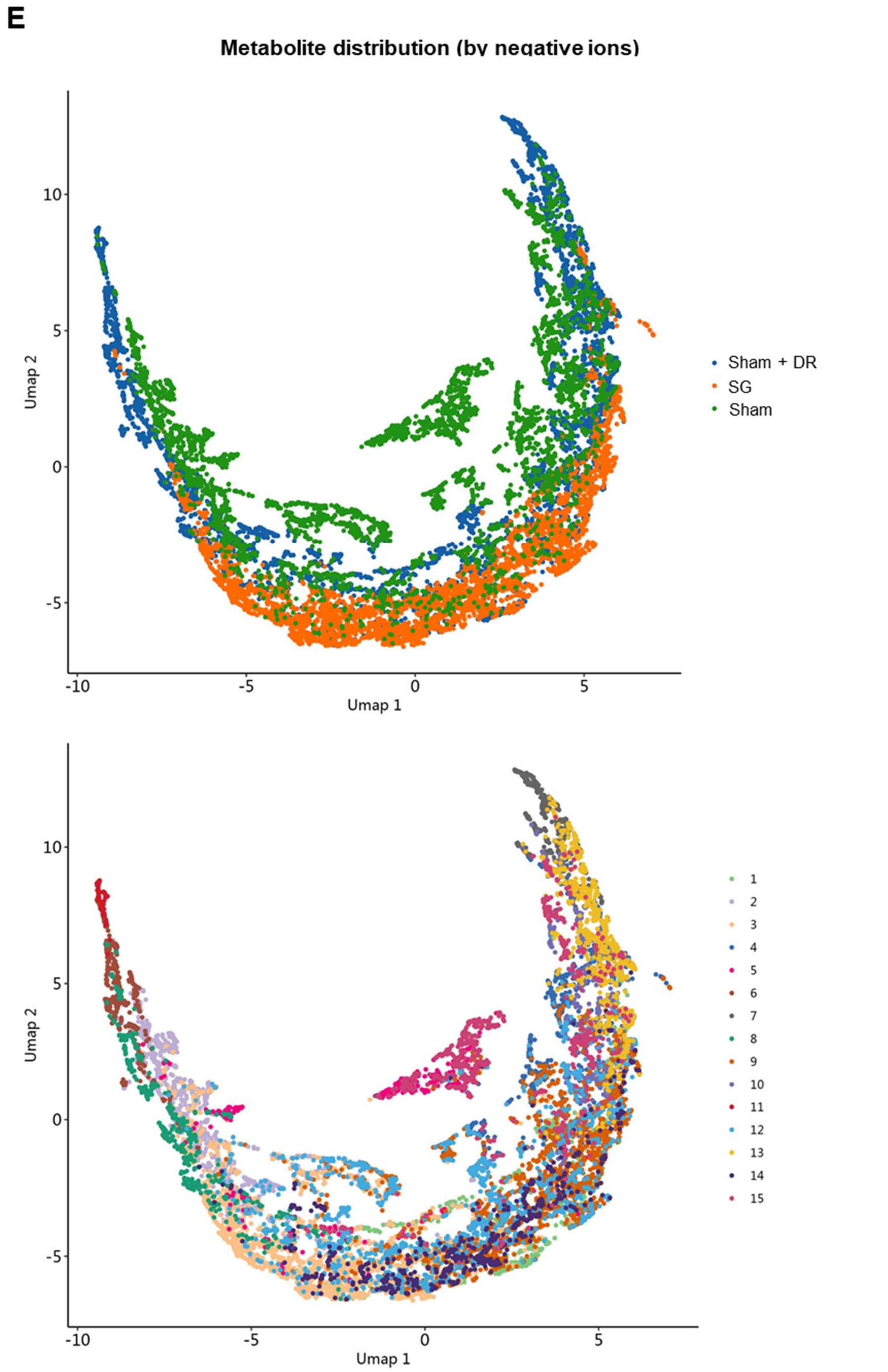

**Figure S6. Metabolomics depicts metabolite profiles and distribution.** Spectrum of mass spectrometry detection by positive and negative ions in **(A)** Sham, **(B)** Sham + DR and **(C)** SG groups. Distribution of metabolites detected by **(D)** positive and **(E)**

negative ions illustrated in UMAPs. Abbreviations: Sham: sham surgery; DR: dietary restriction; SG: sleeve gastrectomy.

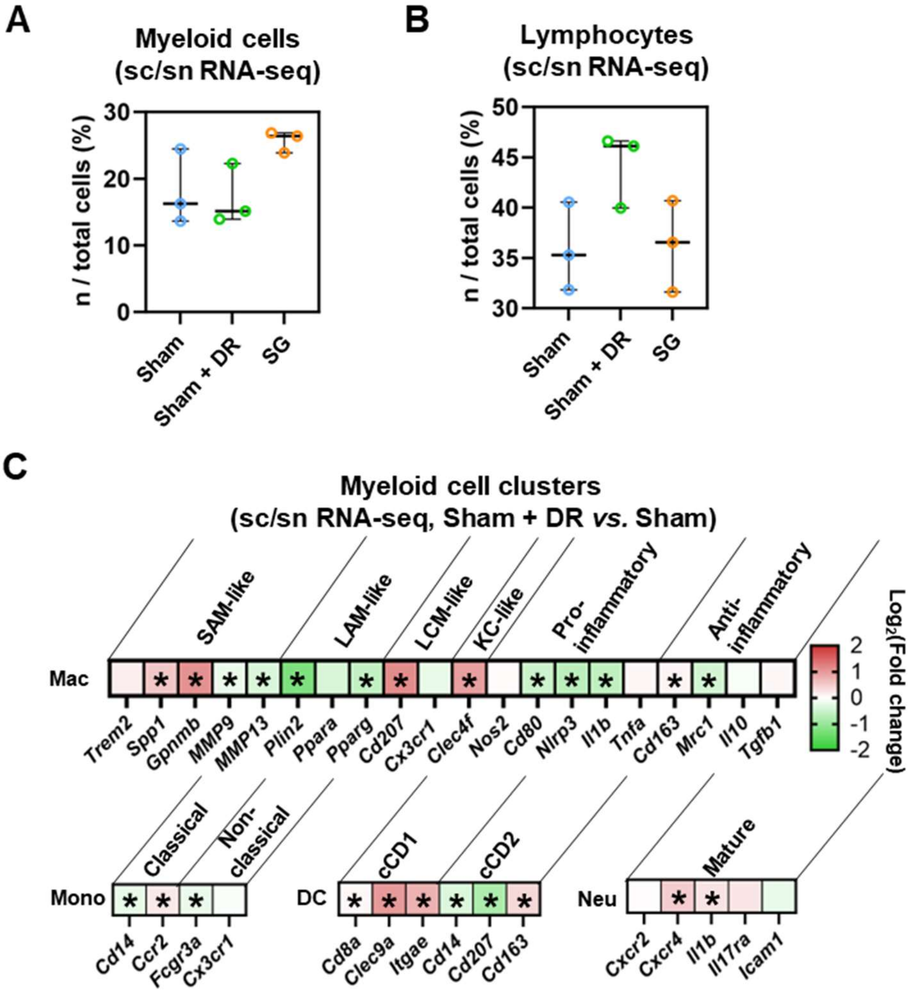

**Figure S7. Assessment of myeloid cells and lymphocytes in rat livers.** Proportions of **(A)** myeloid cells and **(B)** lymphocytes per rat liver sample from sc/sn RNA-seq analysis. Abbreviations: Sham: sham surgery; DR: dietary restriction; SG: sleeve gastrectomy. **(C)** The expression of phenotype-associated markers in myeloid cell clusters were illustrated based on sc/sn RNA-seq data (SG *vs*. Sham). Abbreviations: C: control; Sham/S: sham surgery; DR: dietary restriction; SG: sleeve gastrectomy; KEGG: Kyoto encyclopedia of genes and genomes. sc/sn: single-cell/single-nuclei; Mono: monocytes; Mac: macrophages; DC: dendritic cells; Neu: neutrophils; SAM: scar-associated macrophage; LAM: lipid-associated macrophage; LCM: liver capsular macrophage; KC: Kupffer cell; cDC: conventional DC. The unpaired t-test and one-way ANOVA test were performed. '*' represents '$p < 0.05$' and statistical significance.

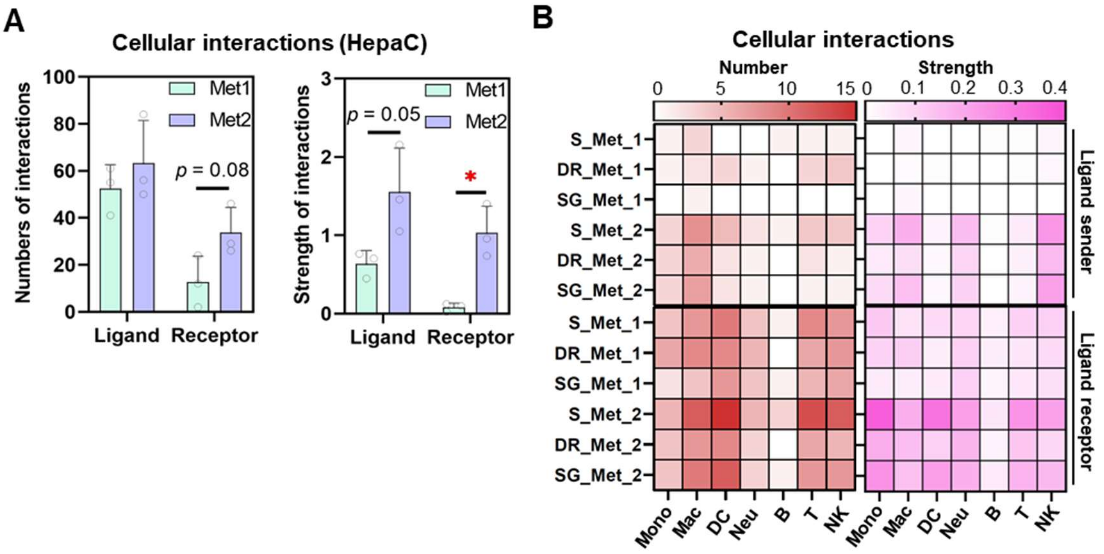

**Figure S8. Cellular interactions in metabolism groups of rat hepatocytes.** Numbers and strength of cellular interactions (ligand-receptor) among hepatocytes (Met 1 and 2) and immune cells (B, T, NK, DCs, macrophages, monocytes and neutrophils) displayed in (A) total values and depicted in matrixes. Abbreviations: Sham: sham surgery; DR: dietary restriction; SG: sleeve gastrectomy; DC: dendritic cells; NK: natural killer cells; B: B lymphocytes; T: T lymphocytes; Neu: neutrophils; Mac: macrophages; Mono: monocytes: HepaC: hepatocytes. The unpaired t-test was performed. '*' represents '$p < 0.05$' and statistical significance.
